# Urban living can rescue Darwin’s finches from the lethal effects of invasive vampire flies

**DOI:** 10.1101/2023.03.06.531275

**Authors:** Sarah A. Knutie, Cynthia N. Webster, Grace J. Vaziri, Lauren Albert, Johanna A. Harvey, Michelle LaRue, Taylor B. Verrett, Alexandria Soldo, Jennifer A.H. Koop, Jaime A. Chaves, Jill L. Wegrzyn

## Abstract

Human activity changes multiple factors in the environment, which can have additive or neutralizing effects on organisms. However, few studies have explored the causal effects of multiple anthropogenic factors, such as urbanization and invasive species, on animals, and the mechanisms that mediate these interactions. This study examines the influence of urbanization on the detrimental effect of invasive avian vampire flies (*Philornis downsi*) on endemic Darwin’s finches in the Galápagos Islands. We experimentally manipulated nest fly abundance in an urban and non-urban area and then characterized nestling health, survival, diet, and gene expression patterns related to host defense. Survival of fumigated nestlings from urban and non-urban nests did not differ significantly. However, sham-fumigated, non-urban nestlings lost more blood and few nestlings survived compared to urban nestlings. Stable isotopic values (δ^15^N) from urban nestling feces were higher than non-urban nestlings, suggesting that urban nestlings are consuming more protein. δ^15^N values correlated negatively with parasite abundance, which suggests that diet might influence host defenses (e.g., tolerance and resistance). Parasitized urban nestlings differentially expressed genes within pathways associated with red blood cell production (tolerance) and pro-inflammatory response (innate immunological resistance), compared to sham-fumigated non-urban nestlings. In contrast, sham-fumigated non-urban nestlings differentially expressed genes within pathways associated with immunoglobulin production (adaptive immunological resistance). Our results suggest that urban nestlings are investing more in pro-inflammatory responses to resist parasites, but also recovering more blood cells to tolerate blood loss. Although non-urban nestlings are mounting an adaptive immune response, it is likely a last effort by the immune system rather than an effective defense against avian vampire flies since few nestlings survived.

## Introduction

Emerging diseases are a major global threat to biodiversity (Daszak et al. 2000, Keesing et al. 2010). Naïve hosts who cannot effectively defend themselves against novel disease-causing parasites may risk population declines or even extinction (van Riper III and van Riper 1986, Frick et al. 2010). However, not all hosts are susceptible to introduced parasites. The fitness of some host species is clearly reduced, while the fitness of other hosts is relatively unaffected. Less affected hosts may alleviate parasite damage with defense mechanisms, such as tolerance and resistance (Read et al. 2008). Tolerance mechanisms, such as tissue repair or recovery of blood loss due to parasites, minimize the damage that parasites cause to the host without reducing parasite fitness (Medzhitov, Schneider, & Soares, 2012; Miller, White, & Boots, 2006; Råberg, Sim, & Read, 2007; Read, Graham, & Råberg, 2008). For example, parents from ectoparasite-infested nests reduce the cost of parasitism by feeding their offspring more than parents from fumigated nests (Christe, Richner, & Oppliger, 1996; Knutie et al., 2016; Tripet & Richner, 1997). Consequently, despite infestation, offspring do not suffer a high cost of parasitism because they are able to compensate for resources lost to the parasites. Alternatively, but not mutually exclusive, resistance, such as the immune response, minimizes the damage that parasites cause to the host by reducing parasite fitness (Read et al., 2008). The presence and effectiveness of host defenses to invasive parasites is highly variable among individuals and populations and can depend on how quickly they can evolve these responses or whether specific environmental factors are available to facilitate them (Feis et al. 2016, McNew et al. 2019).

Humans in urban environments can increase anthropogenic food availability and reliability for animals with the establishment of wild animal feeders or the incomplete disposal of human trash. Due to the high energetic cost of defense mechanisms, only hosts with sufficient food resources, such as in these urban areas, may be able to invest in defenses (Cornet, Bichet, Larcombe, Faivre, & Sorci, 2014; Howick & Lazzaro, 2014; Lochmiller & Deerenberg, 2000; Sheldon & Verhulst, 1996; Sternberg et al., 2012; Svensson, Råberg, Koch, & Hasselquist, 1998; Knutie 2020). Extra nutrients obtained from human-derived food, such as protein, can directly increase the production of immune cells (Coop & Kyriazakis, 2001; Strandin, Babayan, & Forbes, 2018). Consequently, individuals with less effective defenses but better access to resources might be better equipped to resist the negative effects of parasites. Alternatively, human-supplemented food may be of lower nutritional quality, which could decrease the hosts’ health (Catto et al. 2021) and therefore their ability to produce an effective immune response to parasites. Without the development or evolution of resistance and tolerance defenses, host population size can decrease or even go extinct, and this effect might be especially apparent in the context of human-influenced environments (van Riper III and van Riper 1986, Atkinson and Lapointe 2009). Understanding the effects of these complex, non-mutually exclusive interactions is critical because the movement of parasites around the world is only increasing, and the urban ecosystem is one of the few that is rapidly expanding (Birch and Wachter 2011, Zhou et al. 2015, Verrelli et al. 2022).

The effects of urbanization on host defenses against parasites might be particularly pronounced on islands where population sizes are small, genetic diversity is relatively low, and where species have evolved in the absence of introduced parasites (e.g., Hawaiian honeycreepers and malaria). The Galápagos Islands of Ecuador are relatively pristine but face an increasing rate of change as a result of a growing human presence. Ecotourism and the permanent resident human population have grown exponentially in the Galápagos over several decades with nearly 225,000 visiting tourists each year and 30,000 permanent residents (Walsh and Mena 2016). The introduction of novel parasites to the Galápagos is also relatively recent, such as the avian vampire fly (*Philornis downsi*). The adult fly was first collected in the Galápagos in 1964 (Causton et al. 2006), but the first sign of parasitism in nests was in 1997 (Fessl and Tebbich 2002). Adult flies are non-parasitic but lay their eggs in the nests of birds. Once hatched, fly larvae feed on the blood of nestling and brooding female birds (Fessl and Tebbich 2002, Koop et al. 2013). This fly dramatically reduces nestling survival of endemic Darwin’s finches, and in some years, can cause up to 100% mortality of infested nestlings (Koop et al. 2013, O’Connor et al. 2014). However, some endemic Galapagos species, such as the Galapagos mockingbird (*Mimus parvulus*), are able to tolerate the effect of the fly because parents of parasitized nests feed their nestlings more than parents of non-parasitized nests to compensate for energy lost to the flies. Because urban areas offer more consistent food, though likely of lower quality, (Knutie 2020), urban areas could amplify or dampen host defense strategies against this parasitic fly.

This study examines the effect of urbanization on the interactions between Darwin’s finches and invasive avian vampire flies. For our first objective, we investigated whether the effect of the flies on finch nestlings differed in an urban and non-urban area. We experimentally removed vampire flies from or allowed for natural parasitism in the nests of small ground finches (*Geospiza fuliginosa*; a species of Darwin’s finch) in an urban (Puerto Baquerizo Moreno) and non-urban area (Jardin de las Opuntias) on San Cristóbal Island, Galápagos. We then quantified nestling morphometrics, blood loss, and survival in response to parasitism across treatments and sites. Relative diets were compared between urban and non-urban nestlings using stable isotope analyses of feces (δ^13^C, δ^15^N, and C:N). Stable isotope analyses of adult flies were also conducted to determine whether nestling isotope values were reflected in the flies. Because urban birds have better access to resources, which could positively affect immune development, we expect that urban finches will be more resistant to vampire flies than non-urban birds. However, because urban birds prefer human-processed food (De León et al. 2018), which lacks many nutritional qualities required for an active immune system, urban nestlings might not be well-defended against the parasite. Spatial variation of urban parasite abundance was also examined to determine whether parasite load had spatial structure within the urban area or was associated with various environmental features (e.g., restaurants).

Our second objective examined the molecular mechanisms that underlie the defense response to flies in urban and non-urban nestlings. We characterized gene expression in the blood of ∼8 day old nestlings across parasite treatments and locations. Gene expression is the process in which gene information is turned into a functional product, which can affect phenotypes. For example, nestlings in urban areas may have more relative expression of genes in a pathway that affects erythrocyte production, which could account for higher tolerance to flies. Co-expression profiles were also used to compare gene expression with organismal traits of nestlings. If urban nestlings are better defended against flies, we expect to observe increased expression of genes related to innate and adaptive resistance and/or tolerance to blood loss compared to non-urban nestlings. Consequently, urban, sham-fumigated nestlings are expected to have better health and survival as compared to non-urban, sham-fumigated nestlings. Our study is one of the first to provide insight into the mechanisms by which urbanization positively or negatively influences host-parasite interactions.

## Methods

### Study system

We conducted our study between February–May 2019 in an urban and non-urban area in the dry lowland climatic zone of San Cristóbal (557 km^2^) in the Galápagos Islands. The urban area encompasses the only city on San Cristóbal Island, Puerto Baquerizo Moreno (hereon, urban area) (−0.9067715°, −89.6061678°). This capital city is the second largest city in the Galápagos archipelago with a human population of 7,199 (INEC, 2016) and measures 0.79 km^2^ (∼1.2 km by 0.62 km), which includes tourist and residential areas (Harvey et al. 2021). The urban area is almost entirely consumed by human infrastructure, which primarily consists of impermeable concrete or stone surfaces and human built structures, but also includes native plants, such as matazárno (*Piscidia cathagenensi*), Galápagos acacia (*Acacia rorudiana*), and prickly pear cactus (*Opuntia megasperma*), and ornamental non-native plants established by humans.

The non-urban area is in Jardín de las Opuntias (hereon, non-urban area), which is a protected Galápagos National Park site located eight kilometers southeast of the urban area (−0.9491651°, −89.5528454°). Our non-urban study area measures 0.21 km^2^ and covers 1.4 km of the main trail and 0.15 km to each side. The non-urban site does not contain any unnatural, human-built impermeable surfaces, and includes native plants, such as matazárno, acacia, and cacti. The non-urban area receives very low human visitation due to the difficult terrain; however, local residents occasionally transect through the site to access the beach.

Small ground finches on San Cristóbal commonly nest in both the urban and non-urban area, generally between February–May (Harvey et al. 2021). Finches build their nests in matazárno, acacia, and cacti in both locations, but urban finches occasionally nest in human built structures such as gutters and building signs. Finches use coarse grasses and small twigs to build the outer structure of the nest, and finer, softer grasses and plants to construct the inner layer and nest liner. In urban areas, finches frequently incorporate trash and human hair into the construction of their nests (Theodospoulos and Gotanda 2018, Harvey et al. 2021). Finches produce clutches between 1-5 eggs and females incubate the eggs for around 15 days (Harvey et al. 2021). After hatching, nestlings fledge when they are 12-16 days old. Although females are primarily involved in parental care, both males and females feed nestlings via regurgitation.

### Nest parasite manipulation

Both field sites were searched daily or every other day for evidence of nest-building activity by small ground finches. Once a nest was located, it was checked every other day until eggs were laid in the nest. Observations were made primarily through binoculars to minimize nest disturbance. When adults were not at the nest, we used a small camera (Contour LLC), attached to an extendable pole, which transmitted video (via Bluetooth) to an iPhone.

Since vampire flies can lay their eggs during the finches’ egg incubation stage, the nests were assigned to a control (hereon, sham-fumigated) or experimental (hereon, fumigated) treatment after a full clutch of finch eggs was laid. We applied a 1% solution of controlled release permethrin (Permacap: BASF Pest Control Solutions) to experimental nests and water to control nests. Permethrin has been used extensively by Galápagos researchers to experimentally remove avian vampire flies from nests (Fessl et al. 2006, Koop et al. 2013a,b, Knutie et al. 2014, Kleindorfer and Dudaniec 2016, Knutie et al. 2016, McNew et al. 2019, Addesso et al. 2020) and is approved for use by the Galápagos National Park. We treated nests twice with their respective treatments: 1) during the egg stage, and 2) when nestlings hatched. During the first treatment, 3 mL of permethrin or water was injected beneath the nest liner with a sterile blunt syringe. During the second treatment, the nest liner and nestlings were removed from the nest and the permethrin or water was applied by spraying (10 times) into the base of the nest with a travel-sized spray bottle. The treatments were applied below the nest liner to ensure that the eggs and nestlings did not directly contact the permethrin or water. While hatchlings were outside the nest during nest treatment, they were checked for signs of vampire fly infestation (black or enlarged nares, or blood on legs, wings, or feather pores). The nest liner and nestlings were then returned to the nest, Julian hatch day was recorded, and a GPS coordinate was taken for each nest.

### Nestling health and sample collection

We returned to the nest when nestlings were 7-8 days old to measure their body mass (to the nearest 0.1 g) with a portable digital scale balance (Ohaus CL2000) and morphometrics, such as tarsus, bill length, bill width, and bill depth (to the 0.01 mm) with analog dial calipers from Avinet. During this visit we opportunistically collected fecal samples from the nestlings. Briefly, a nestling was held approximately 10 cm over a sterile plastic weighboat until it defecated (<10 seconds). We then transferred the fecal sac into a sterile 2 mL tube. The fecal sample was transferred to a 2mL tube and kept at 4°C in a portable insulin refrigerator until we returned from the field. Samples were stored in a −20°C freezer while in the Galápagos and then transferred to the University of Connecticut where they were stored at −80°C until processed for stable isotope analysis.

For up to three nestlings per nest, we also collected a small blood sample (<20 µL) from the brachial vein using a 30-gauge sterile needle and heparinized capillary tube. Since nestlings are being fed upon by a hematophagic ectoparasite, we wanted to collect the smallest blood sample possible from each nestling. Therefore, we used a blood sample from each nestling within a nest for a different assay. One sample was used to quantify hemoglobin levels (g/dL) with a HemoCue® HB +201 portable analyzer (Hemocue America, USA). The second sample was used to quantify glucose levels (mg/dL) with a glucometer (OneTouch, USA). The third sample of whole blood was preserved in 180 µL of RNAlater; this preserved blood was kept at 4°C in a portable insulin refrigerator until we returned from the field. The sample was then vortexed and stored at 4°C for 24 h, according to manufacturer’s protocol, before being placed in a −20°C freezer while in the Galápagos (up to two months). The samples were then transported on ice packs (20 h) to the University of Connecticut where they were stored for two months at −80°C until extracted for RNA sequencing.

We banded nestlings with an individually numbered metal band and a unique combination of three colored bands. When nestlings were approximately 12 days old, we observed the nest every two days with binoculars from a distance of approximately 5 m (to prevent premature fledging). Successful fledging was confirmed by identifying individual birds once they left the nest.

### Quantifying parasite abundance

Within two days of all nestling birds either fledging or dying, the nest was collected and placed in a sealed, gallon-sized plastic bag. Nests were transported from the field and dissected by hand to collect all stages of vampire flies present in the nest within 8 h. All larvae (1^st^, 2^nd^, and 3^rd^ instars), pupae, and pupal cases were identified and counted to determine total parasite abundance for each nest. The length and width (0.01 mm) of up to ten pupae were haphazardly measured with digital calipers. These measurements were used to calculate pupal volume (V = π*[0.5*width]^2^*length). Third instar larvae were placed in ventilated 50 mL Falcon tubes with their home nest material until pupation. Pupae were also placed in 50 mL Falcon tubes (without material) until they eclosed. Once they eclosed, up to ten adult flies were collected and preserved in 95% ethanol for stable isotope analysis (see below for methods).

### Stable isotope analyses

We quantified δ^13^C (^13^C:^12^C), δ^15^N (^15^N:^14^N), and the carbon to nitrogen (C:N) ratio of finch feces and adult fly bodies. δ^13^C helps explain the differences in C4 vs. C3 plants consumed, δ^15^N helps explain the amount of dietary protein consumed (and infers trophic level), and C:N helps explain the relative lipid consumption. Stable isotope analysis was used for flies to examine whether nutritional differences were detectable in the vampire flies parasitizing the finches. Feces were dried in an oven at 60°C for 24 h. Fly samples were rinsed three times in a 2:1 (vol/vol) chloroform/methanol mixture to remove surface oils, then rinsed with sterile water, and dried at 60°C for 24 h. After drying, samples were ground to a fine powder using a mortar and pestle. Up to 1 mg of each dried, homogenized sample was weighed into a tin capsule (Costech Analytical Technologies, Inc., USA). Capsules were folded, and placed into a 96-well plate, then sent to the University of New Mexico Center for Stable Isotope Ratio Analysis (SIRA). The samples were run as duplicates on a Thermo Delta V mass spectrometer connected with a Costech 4010 Elemental Analyzer.

### Effect of location and landmarks on parasite abundance and fledging success

To explore whether location or landmarks (i.e. food markets, bakeries, benches and restaurants) are associated with parasite abundance and fledging success, we conducted an optimized hotspot analysis in ArcGIS 10.8.1 (ESRI 2020). First, a minimum convex polygon (MCP) was created to serve as our study area for spatial analysis and included all landmarks within the urban area. Nesting hotspots were then determined based exclusively on spatial location of nests (i.e., clusters of finch nests rather than any other features on the landscape), which resulted in a grid of cells that were characterized as hotspots or not within the MCP-derived study area. To learn about the characteristics of these nests, the Near Tool was used to identify and calculate the distance to the nearest landmark for all urban nest locations that were within the nesting hotspots. Specifically, we were interested in all landmarks associated with food, which was 190 records of 303 landmarks available. We then extracted the locations of nests within hotspot locations and reviewed landmark types associated with these nests. Finally, we conducted hotspot analyses based on parasite load (i.e., is there a hotspot of nest flies?) and also based on fledging (i.e., is there a hotspot of fledging success?). The hotspot analysis uses the Getis Ord-Gi statistic (Getis and Ord 1992), which determines spatial clusters with either high or low values for the statistic (Fisher and Getis 2010), corresponding to hot or cool spots, respectively.

### RNA extractions, sequencing, and bioinformatics

Total RNA was extracted from 20 µL of peripheral whole blood using a modified Tri-Reagent (Ambion, Invitrogen, USA) and Direct-zol RNA Miniprep Plus Kit (Zymo Research, USA) protocol (Harvey and Knutie 2022). The samples were incubated at room temperature for 2 min and then centrifuged for 1 min at 8,000 x g to lightly pellet the blood. Preservation fluid (RNAlater, Ambion, Invitrogen, USA) was pipetted off leaving no more than ∼15 µL of preservative and 500 µL of Tri-Reagent were added along with a sterile 5 mm stainless steel bead (Thomson, USA). The samples were then vortexed for 30 seconds before adding an additional 500 µL of Tri-Reagent. The sample was then vortexed for 10 min at room temperature. The phase separation portion of the Tri-Reagent protocol was then followed, and the upper aqueous phase (500 µL) was transferred to new microcentrifuge tubes. We then followed the manufacturer’s protocol for the Direct-zol RNA Kit beginning with the RNA purification step. We eluted total RNA using 50 µL of RNA/DNA free water. We used a 4200 TapeStation and High Sensitivity RNA ScreenTape assays (Agilent, USA) to quantify total RNA concentration (ng/µL) and RNA integrity numbers (RIN^e^) (Schroeder et al. 2006). RNA extracts were then stored at −80 °C until sequencing. The mean RNA concentration was 74.62 ng/µL (range: 15.2-246.0) and mean RIN^e^ was 9.33 (range: 7.8-10), with all samples above the minimum RIN^e^ cutoff (= 7.0) for successful sequencing.

A poly-A tail binding bead-based approach was used to reduce ribosomal RNA contamination. First- and second-strand cDNA was synthesized using the Illumina TruSeq Stranded mRNA Sample preparation kit and dual indexing was used to multiplex sequencing of samples. Library quality was assessed on the Agilent Tapestation D1000 DNA High Sensitivity assay and quantified using the Qubit 3.0 High Sensitivity dsDNA assay to ensure equimolar pooling. A total of 42 libraries were sequenced across two (75bp PE) Illumina NextSeq500 High Output sequencing runs.

Quality control was applied to paired-end libraries via Trimmomatic (v.0.39) (minimum quality score 20 and minimum length 45bp) (Bolger et al. 2014). The trimmed reads were aligned to the reference genome, *Geospiza fortis* (GeoFor_1.0, INSDC Assembly GCA_000277835.1, Jul 2012), utilizing HISAT2 (v.2.2.1) (Kim et al. 2015). Read counts were extracted from each alignment file via HTSeq (v.0.13.5) with the published reference annotation (Anders et al. 2015).

Read counts and corresponding treatment/location assignments were imported into RStudio (Bioconductor), and DESeq2 (v.1.32.0) was used to identify differentially expressed genes among nestling groups (Soneson et al. 2015; Love et al. 2014). The factorial design with an interaction term (∼site + treatment + site:treatment) compared urban and non-urban sites, and sham-fumigated and fumigated treatments (Fig. S2). *P*-values derived from the Wald test were corrected for multiple testing using the Benjamini and Hochberg method (Love et al. 2014). We selected genes with an adjusted *P*-value less than 0.1. Initial functional annotations were imported via biomaRt (v.2.48.3) with Ensembl (bTaeGut1_v1.p) (Durinck et al. 2009). An enrichment and depletion (two-sided hypergeometric test) *immune system process* Gene Ontology (GO) enrichment analysis was performed on differentially expressed genes of sham-fumigated urban and non-urban nestlings to predict potential immune function pathways. *Taeniopygia guttata* (zebra finch) GO annotations were sourced from UniProt GOA and referenced within ClueGO (v.2.5.9) and CluePedia (v.1.5.9), plug-ins for Cytoscape (v.3.8.2) (Bindea et al. 2009; Bindea et al. 2013; Shannon et al. 2003). Terms with a *P*-value < 0.1 were considered significantly enriched.

A co-expression analysis was conducted via WGCNA (v.1.71) in RStudio (Langfelder and Horvath 2008). Binary and quantitative trait information was imported along with normalized RNA-Seq counts. A soft-thresholding power of 6 was chosen based on the criterion of approximate scale-free topology and sample size, and eigengene significance was calculated for each module. Within Cytoscape, a ClueGO network (referencing zebra finch) was constructed to visualize key drivers in the context of immune response.

### Statistical analyses

Statistical analyses of field data (i.e., non-gene expression data) were conducted in R (2021, v.1.4.1103) and figures were created in Prism (2021, v.9.2.0). For dependent variables with continuous data, we used a Shapiro-Wilk test for normality then chose the appropriate distribution for the analysis. Since mass and tarsus were highly correlated (R^2^ = 0.74, *P* < 0.0001), we calculated the scaled mass index, which is a standardized metric of body condition (Peig and Green 2009). Bill surface area was also calculated from bill length, width, and depth using a modified equation for the surface area of a cone (Greenberg et al. 2012).

Linear mixed effects models (LMMs) with nest as a random effect were used to analyze the effect of location on parasite volume, effect of parasite intensity on parasite size, and effect of Julian hatch day on parasite size. LMMs were used to determine the effect of site, treatment, and their interaction on scaled mass index, bill size, hemoglobin levels, and glucose levels, with nest as a random effect. Age and Julian hatch day were included as covariates when they contributed significantly (*P* < 0.05) to the model (age only: bill size; Julian hatch day only: glucose; neither: scaled mass index, hemoglobin levels). LMMs were also used to determine the effect of site and sample type (nestling feces vs. fly) on δ^15^N, δ^14^C, and C:N, with sample replicate as a random effect.

Generalized linear models (GLMs) were used to analyze the effect of location on parasite abundance (Poisson distribution), the effect of location on Julian hatch day (Gamma distribution), the effect of location and Julian hatch day on brood size (Gamma distribution), and effect of Julian hatch day on parasite abundance (Poisson). GLMs were also used to determine the effect of site, treatment, and their interaction on fledging age for the nest (log transformed; Gaussian). A GLM with binomial errors for proportional data (i.e., a logistic regression) was used to determine the effect of site, treatment, and their interaction on fledging success with Julian hatch day as a covariate.

Analyses were conducted with the lmer function (LMMs) and glm function (GLMs) using the lme4 package in R (Bates et al. 2015). Probability values were calculated using log-likelihood ratio tests using the Anova function in the car package (Fox and Weisberg 2019). For one-way and two-way ANOVAs, we used type II and type III sum of squares, respectively.

## Results

### Effect of parasite treatment and urbanization on parasite load

Urban and non-urban finches initiated breeding in early February 2019. The first urban nests hatched approximately seven days before the first non-urban nest. Although Julian hatch day was, on average, earlier for urban nests (mean ± SE: 86.24 ± 3.99 days) compared to non-urban nests (94.17 ± 4.07 days), Julian hatch day did not differ significantly between sites (χ^2^ = 1.85, *P* = 0.17). Furthermore, site, Julian hatch day, and their interaction did not significantly affect brood size (urban: 2.66 ± 0.18 nestlings, *n* = 38 nests; non-urban: 2.87 ± 0.17, *n* = 30 nests; site: χ^2^ = 0.44, *P* = 0.51; hatch day: χ^2^ = 0.68, *P* = 0.41; interaction: χ^2^ = 1.12, *P* = 0.29). Eight nests were classified as depredated, either due to direct evidence of depredation (e.g., body parts found) or because they were likely depredated (e.g., all nestlings disappeared over a short period of time and prior to the typical fledging window; Table 1).

**Table 1.**
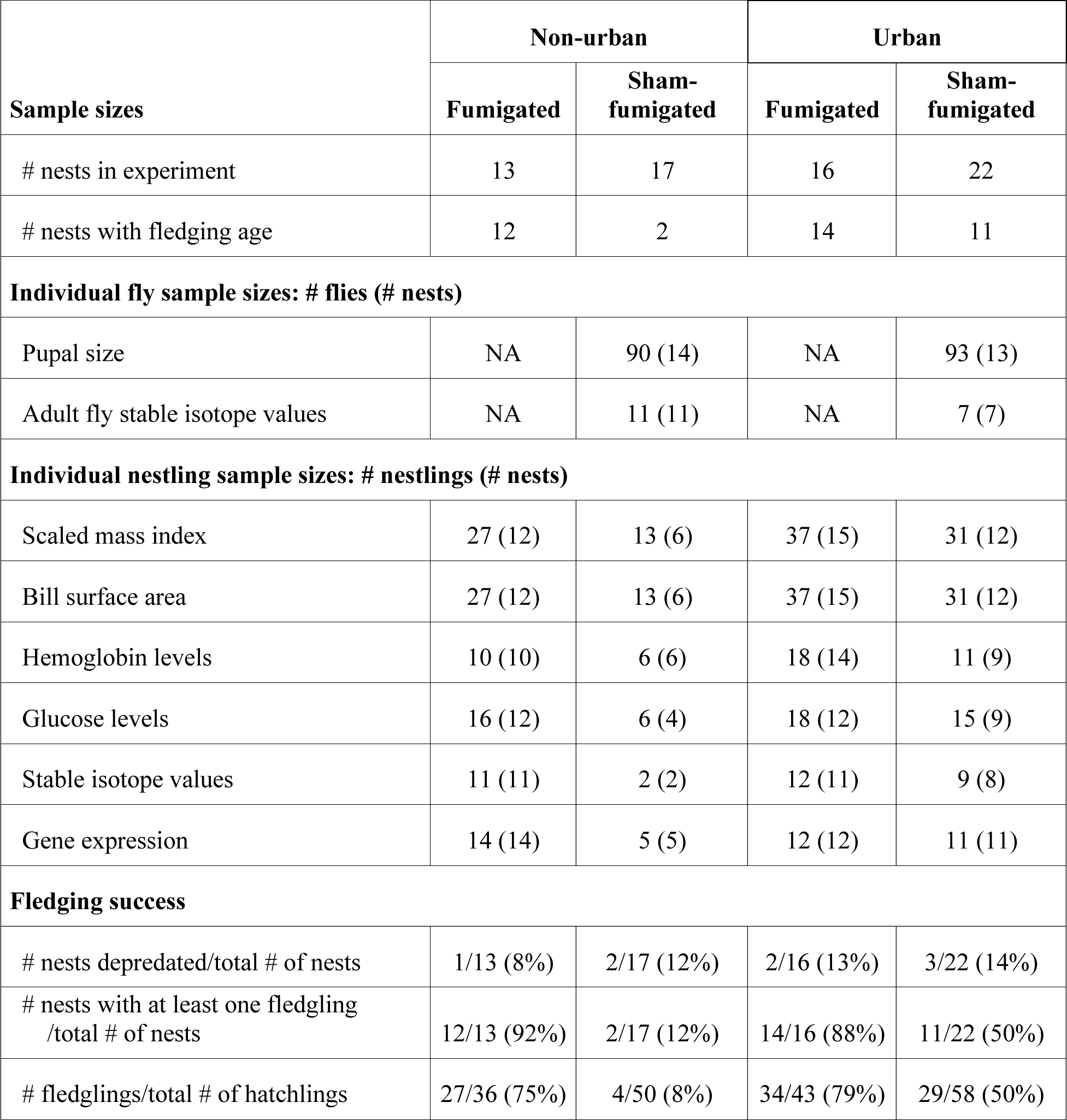
Sample sizes for the effect of location (non-urban or urban) and parasite treatment (fumigated or sham-fumigated) on nest, parasite, and nestling variables.

The experimental treatment of nests with permethrin was nearly 100% effective at removing flies (Fig. 1A; Table 1); only one permethrin-treated urban nest had eight flies after the treatment. Water-treated nests were naturally parasitized by 27.00 (± 5.20) flies in the non-urban area and 15.95 (± 3.15) flies in the urban area (Fig. 1A; Table 1). One water-treated nest disappeared due to predation after the nestlings were banded and therefore we could not quantify parasitism. Fourteen of 17 (82.35%) non-urban nests and 17 of 21 (80.95%) urban nests contained at least one parasite. Within the sham-fumigated treatment, non-urban nests had significantly fewer flies than urban nests (χ^2^ = 54.49, *P* < 0.0001). Julian hatch day did not significantly predict nest parasite abundance (χ^2^ = 1.58, *P* = 0.21).

**Fig. 1.**
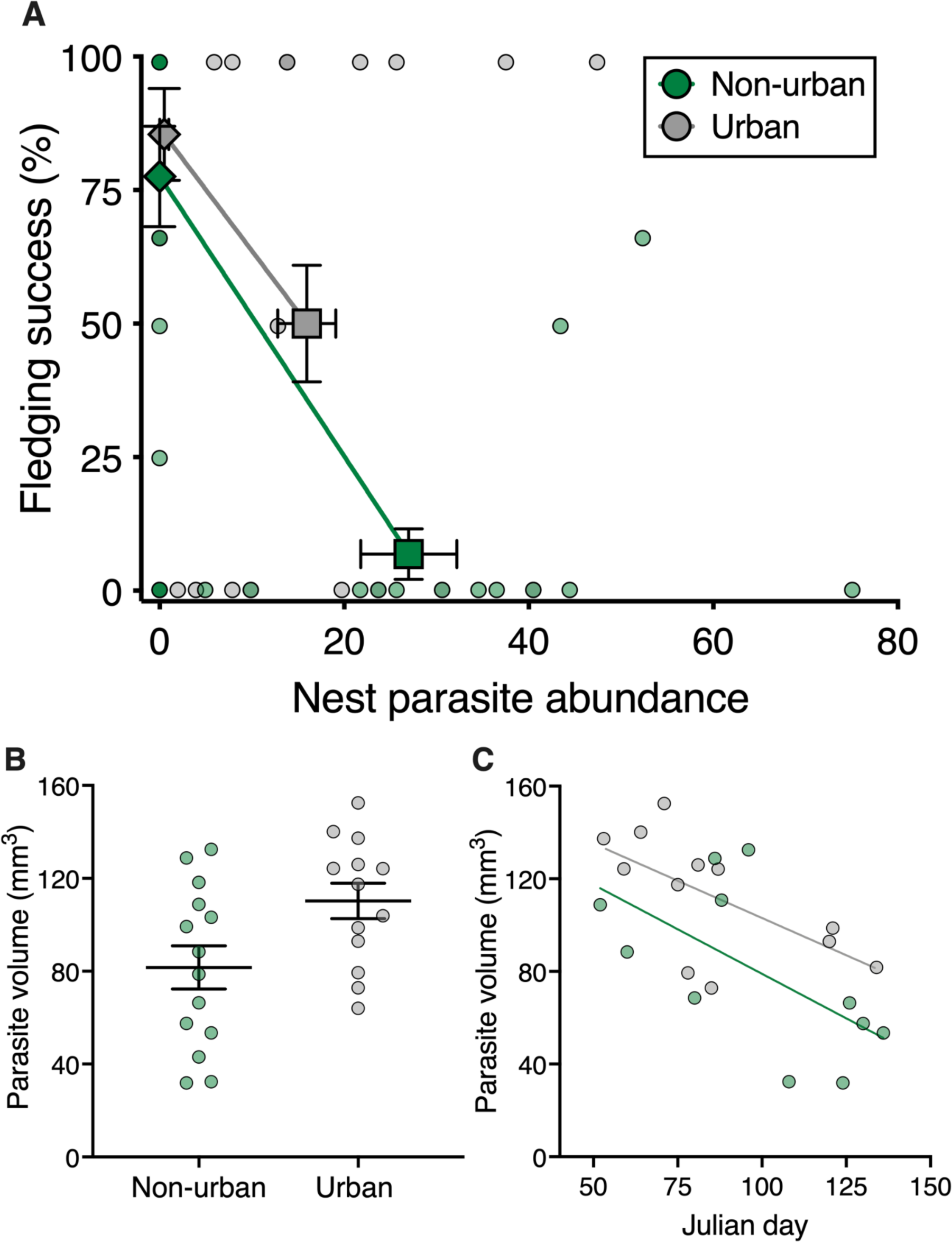
A) The influence of urbanization on mean (± SE) nest parasite abundance (number of flies per nest) and % fledging success. Fledging success did not differ for fumigated nests in the urban and non-urban area. In sham-fumigated nests, urban nests had, on average, fewer flies and higher fledging success compared to non-urban nests. Each point represents an individual nest and darker points indicate higher sample sizes. Diamonds represent the fumigated treatment and squares represent the sham-fumigated treatment. B) Pupal volume (mm^3^) was larger in the urban area compared to the non-urban area. Each point represents a mean (± SE) volume for a nest. C) Parasite volume decreased throughout the nesting season in both locations. Means and standard error bars are represented in panels A and B.

Parasite volume was larger in urban nests compared to non-urban nests (Fig. 1B; χ^2^ = 5.85, *P* = 0.02; Table 1). Urban fly pupae were, on average, 26% larger than non-urban fly pupae(urban: 100.30 ± 7.62 mm^3^, non-urban: 81.61 ± 9.29 mm^3^). This difference was not related to competition among flies within the nest because parasite intensity was not related to parasite size (χ^2^ = 0.01, *P* = 0.97). Rather, the nest hatch day predicted parasite size, with earlier nests having larger flies compared to later nests (Fig. 1C; χ^2^ = 11.01, *P* < 0.001).

### Effect of location and landmarks on nesting and parasite abundance

Of all urban nests (sham-fumigated and fumigated), one nesting hotspot was identified within the urban area (*n* = 37 total nests; *n* =19 hotspot nests), which was at the airport (Fig. S1). The hotspot grid resulted in 14 cells (78 m x 78 m) with 90% confidence of being a hotspot, with *Z*-scores ranging between 2.90 and 6.02. The remaining cells in the study area were not identified as significant nesting hot- or cool-spots. Of the 19 sham-fumigated nests within the nesting hotspot, seven contained flies, and all but one of these was located near a bench. Hotspots were not identified for fledging success, nor did we identify parasite hotspots among sham-fumigated nests.

### Effect of parasitism and urbanization on nestling health

Nestlings from sham-fumigated nests had, on average, lower body condition (i.e., scaled mass index) than nestlings from fumigated nests (Table S1; χ^2^ = 4.01, *P* = 0.045). Site and the interaction between site and treatment did not significantly affect body condition (site: χ^2^ = 0.36, *P* = 0.55; interaction: χ^2^ = 2.20, *P* = 0.14). Treatment and site did not significantly affect bill surface area (site: χ^2^ = 0.13, *P* = 0.72; treatment: χ^2^ = 0.71, *P* = 0.40). The interaction between treatment and site had a marginally non-significant effect on bill surface area (χ^2^ = 3.09, *P* = 0.08); fumigated nestlings from non-urban areas had larger bill surface area than nestlings from urban areas but the bill surface area for sham-fumigated nestlings from urban and non-urban nests did not significantly differ. Most nestlings from non-urban, sham-fumigated nests died before they could be measured; only 13 (of 44) non-urban nestlings from six sham-fumigated nests survived to be measured.

Site alone did not significantly affect nestling hemoglobin levels (Fig. 2; Table S1; χ^2^ = 1.06, *P* = 0.30). Overall, sham-fumigated nestlings had lower hemoglobin levels than fumigated nestlings (χ^2^ = 18.82, *P* < 0.0001). Furthermore, the interaction between treatment and site affected hemoglobin levels (χ^2^ = 6.16, *P* = 0.01), with nestlings from sham-fumigated nests in the non-urban area having, on average, 32% lower levels than nestlings from the other treatments (i.e., sham-fumigated urban, fumigated urban, and fumigated non-urban nestlings).

**Fig. 2.**
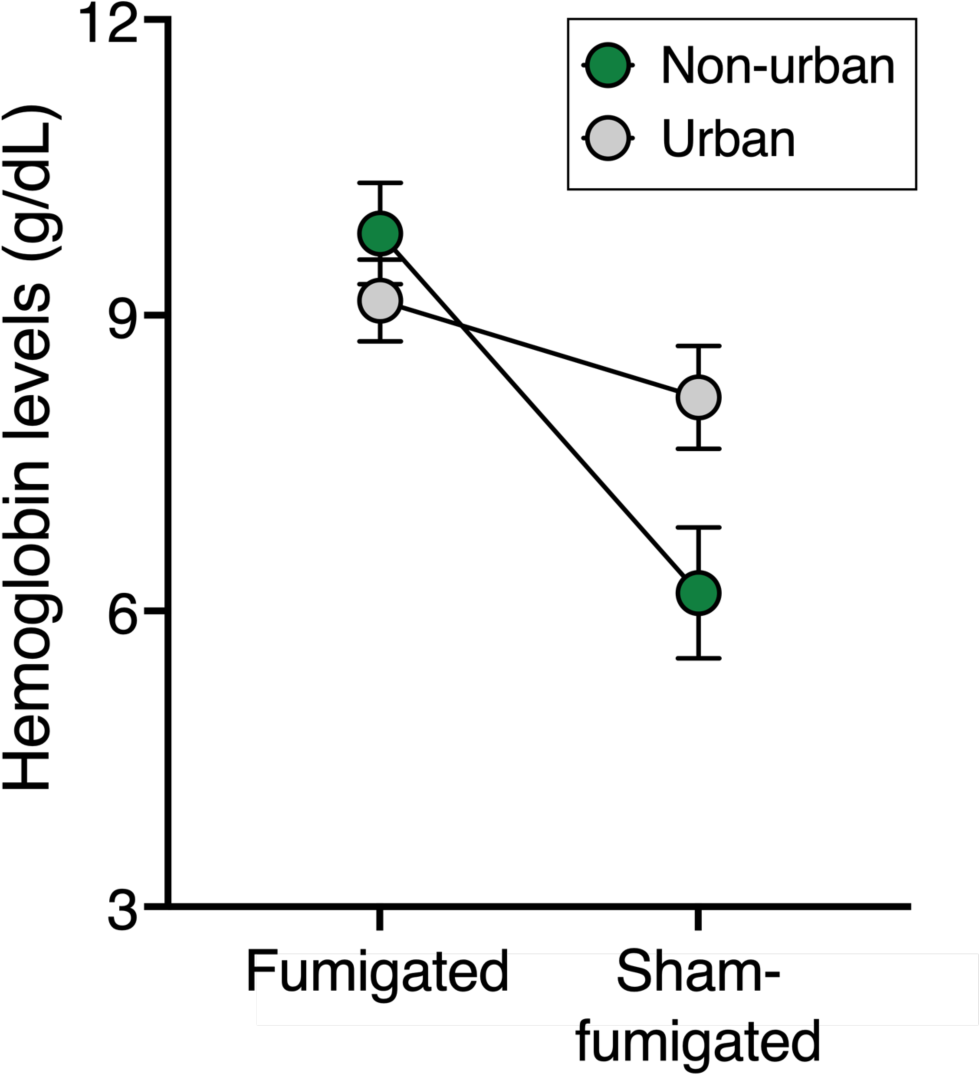
Effect of parasitism and urbanization on mean (± SE) blood loss (hemoglobin) in nestling finches. Nestlings from sham-fumigated nests had lower hemoglobin levels than nestlings from fumigated nests. Non-urban nestlings from sham-fumigated nests had lower hemoglobin levels than nestlings from the other treatments and locations.

Overall, nestlings from fumigated nests had higher glucose levels than nestlings from sham-fumigated nests (Table S1; χ^2^ = 9.75, *P* = 0.002) and non-urban nestlings had higher glucose levels than urban nestlings (χ^2^ = 6.87, *P* = 0.009). Additionally, the interaction between treatment and site had an effect on nestling glucose levels (χ^2^ = 9.34, *P* = 0.002), with urban nestlings from fumigated and sham-fumigated nests maintaining similar levels but non-urban nestlings from fumigated nests having higher levels than sham-fumigated nestlings. Nestlings from sham-fumigated nests in the non-urban area had, on average, 26% lower glucose levels than nestlings from the other treatments (i.e., sham-fumigated urban, fumigated urban, and fumigated non-urban nestlings).

Fledging success of urban and non-urban nestlings did not differ significantly (χ^2^ = 0.67, *P* = 0.41), but overall, parasitism reduced fledging success (Fig. 1A; Table S1; χ^2^ = 44.30, *P* < 0.0001). The interaction between treatment and site affected fledging success (χ^2^ = 8.28, *P* = 0.004); survival of fumigated nestlings did not differ between sites but survival of sham-fumigated nestlings was lower in non-urban areas, compared to urban areas (Table 1). Age at fledging did not differ significantly between treatments (χ^2^ = 0.20, *P* = 0.66) but differed between locations (χ^2^ = 3.71, *P* = 0.05); urban nestlings left the nest approximately one day later than non-urban nestlings. The interaction between treatment and site did not significantly affect fledging age (χ^2^ = 0.10, *P* = 0.75).

### Effect of urbanization on finch and parasite diet

Flies were enriched in δ^15^N compared to nestling feces (χ^2^ = 164.94, *P* < 0.0001) and all urban samples were enriched in δ^15^N compared to non-urban samples (χ^2^ = 108.13, *P* < 0.0001; Fig. 3A). However, the interaction between sample type and site did not significantly affect δ^15^N (χ^2^ = 0.00, *P* = 0.99). Nest parasite abundance correlated negatively with δ^15^N values (χ^2^ = 5.69, *P* = 0.02; Fig. 3B). Nestling feces were enriched in δ^13^C compared to flies (χ^2^ = 11.21, *P* = 0.0008). However, site or the interaction between sample type and site did not affect δ^13^C (site: χ^2^ = 0.43, *P* = 0.51; interaction: χ^2^ = 1.16, *P* = 0.28). Nestling feces had a higher carbon to nitrogen ratio (C:N), compared to flies (χ^2^ = 21.60, *P* < 0.0001). However, site and the interaction between sample type and site did not affect C:N (site: χ^2^ = 2.04, *P* = 0.15; interaction: χ^2^ = 0.74, *P* = 0.39).

**Fig. 3.**
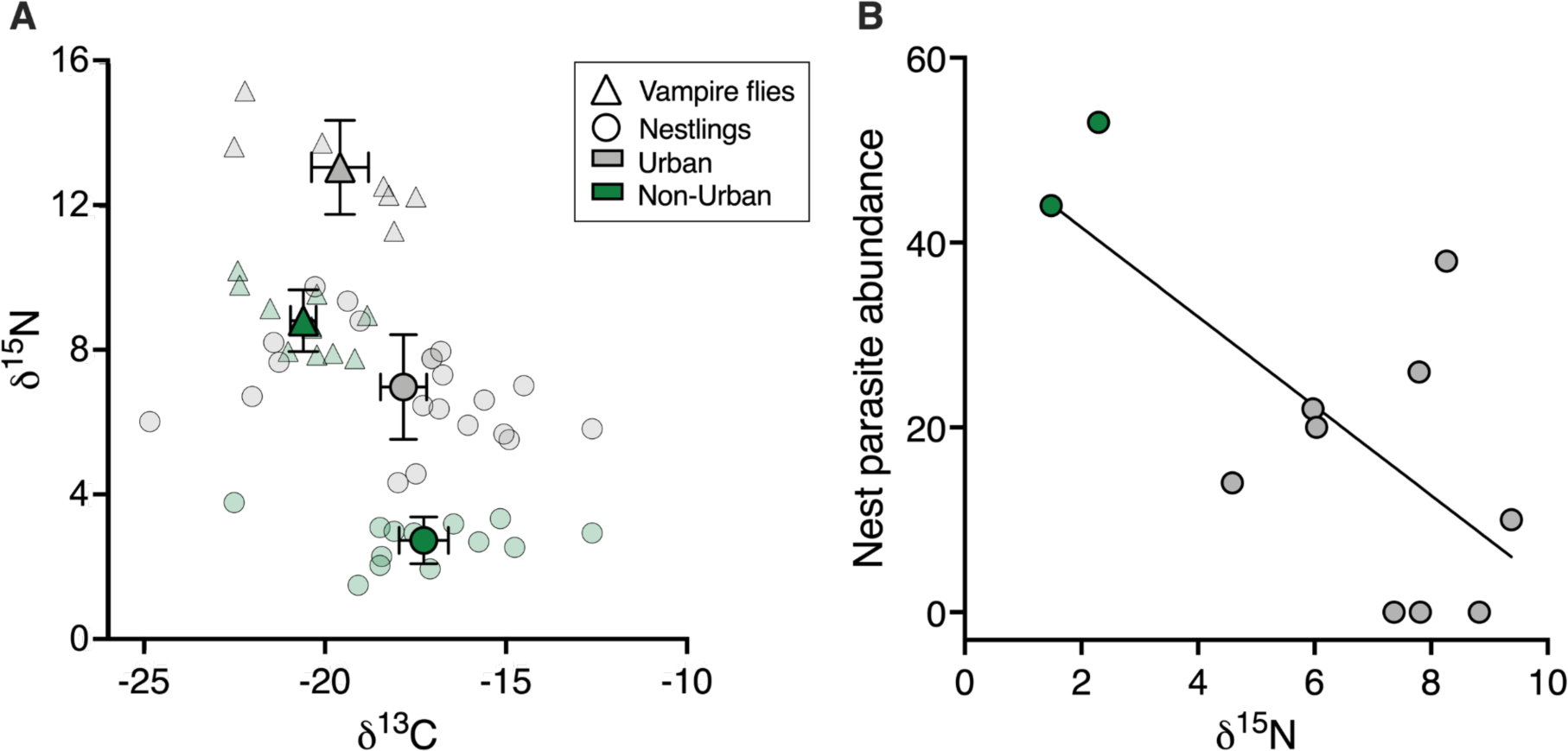
A) Mean (± SE) δ^15^N and δ^13^C values of nestlings and flies in the urban and non-urban area. Urban individuals were enriched with δ^15^N compared to non-urban individuals. Overall, flies were enriched with δ^15^N compared to nestlings and nestlings were enriched with δ^13^C compared to flies. B) Nest parasite abundance (number of flies per nest) was correlated negatively with δ^15^N values. Each point represents an individual.

### Sequencing, Quality Control, and Alignment

In total, 42 paired-end nestling libraries were constructed, including 11 urban sham-fumigated, five non-urban sham-fumigated, 14 urban fumigated, and 12 non-urban sham-fumigated nestlings (1 nestling per nest). Altogether, the paired-reads post-QC ranged between 15.86M and 42.16M and alignment rates against the *Geospiza fortis* genome ranged from 80.74% to 91.81% (Table S2).

### Differentially Expressed Genes

A pairwise differential expression analysis observing two sites (urban and non-urban) and two treatments (fumigated and sham-fumigated), with an interaction, was conducted (File S1). A total of 5,123 genes (*P-adj* < 0.1) were differentially expressed in sham-fumigated finches across the urban (up-regulated) and non-urban (down-regulated) sites, of which, 2,521 were up-regulated and 2,602 were down-regulated. Altogether, 57 genes demonstrated strong expression patterns (± 4 fold change), including frizzled class receptor 10 FZD10 (+16.20 fold change), a primary receptor for Wnt signaling, leucine rich repeat and Ig domain containing 3 LINGO3 (+5.35 fold change), which has been shown to regulate mucosal tissue regeneration in humans and promote wound healing, adenosine deaminase ADA (−8.17 fold change), associated with hemolytic anemia, and hepatic leukemia factor HLF (−5.64 fold change), known to influence the renewal of hematopoietic stem cells (Wang et al. 2016; Zullo et al. 2021; Chen & Mitchell 1994; Komorowska et al. 2017). In fumigated finches, only two genes were up-regulated in the urban site, bisphosphoglycerate mutase BPGM (+3.24 fold change), a regulator of erythrocyte metabolism and hemoglobin in red blood cells, and zinc finger protein GLIS1 (+3.40 fold change), which has been significantly associated with bill length (Xu et al. 2020; Lundregan et al. 2018). There were no significantly down-regulated genes.

When comparing sham-fumigated versus fumigated finches in the urban site, a total of 768 genes were differentially expressed, with 570 being up-regulated in sham-fumigated birds, and 198 down-regulated. The vast majority of genes showed moderate-to-low fold-change patterns, with only three exhibiting strong expression patterns: inorganic pyrophosphate transport regulator ANKH (+4.43 fold change), prokineticin 2 PROK2 (+4.38 fold change) and plexin A2 PLXNA2 (−4.08 fold change).

By comparison, sham-fumigated versus fumigated finches in the non-urban site yielded 7,064 differentially expressed genes, 3,709 of which were up-regulated in sham-fumigated birds, and 3,355 down-regulated. Again, the majority of genes exhibited moderate-to-low fold-change, with only 314 displaying strong expression patterns. This included an assembly factor for spindle microtubules ASPM (+6.34 fold change), which has an apparent role in neurogenesis and neuronal development (Nam et al. 2010) (Fig. S3-S4).

### Gene enrichment related to host defense mechanisms

Gene enrichment of sham-fumigated nestlings from the urban and non-urban site was compared to identify potential host defense mechanisms to explain the expression patterns observed. Analysis of enriched *immune system process* Gene Ontology (GO) terms revealed 21 significant (*P <* 0.05) up-regulated terms (urban) and 12 significant down-regulated terms (non-urban) categorized by resistance (adaptive and innate) and tolerance (File S2).

First, we examined enrichment patterns related to *adaptive immunological resistance* (Fig. 4A). The urban sham-fumigated nestlings exhibited strong enrichment of lymphocyte differentiation (*P* < 0.02) and T cell pathways related to regulation of CD4-positive, alpha-beta T cell activation (*P* < 0.01) and NK T cell differentiation (*P* < 0.05). By comparison, non-urban sham-fumigated nestlings yielded 21 significant adaptive immunity terms. Of those, ten were related to Ig antibodies, including positive regulation of isotype switching to IgG isotypes (*P <* 0.001) and somatic hypermutation of immunoglobulin genes (*P* < 0.001). The rest related to B cell proliferation (*P* < 0.01) and activation (*P* < 0.05), leukocyte differentiation (*P <* 0.04) and proliferation (*P* < 0.05), regulation of antigen receptor-mediated signaling pathways (*P* < 0.04) and germinal center formation (*P* < 0.01).

**Fig. 4.**
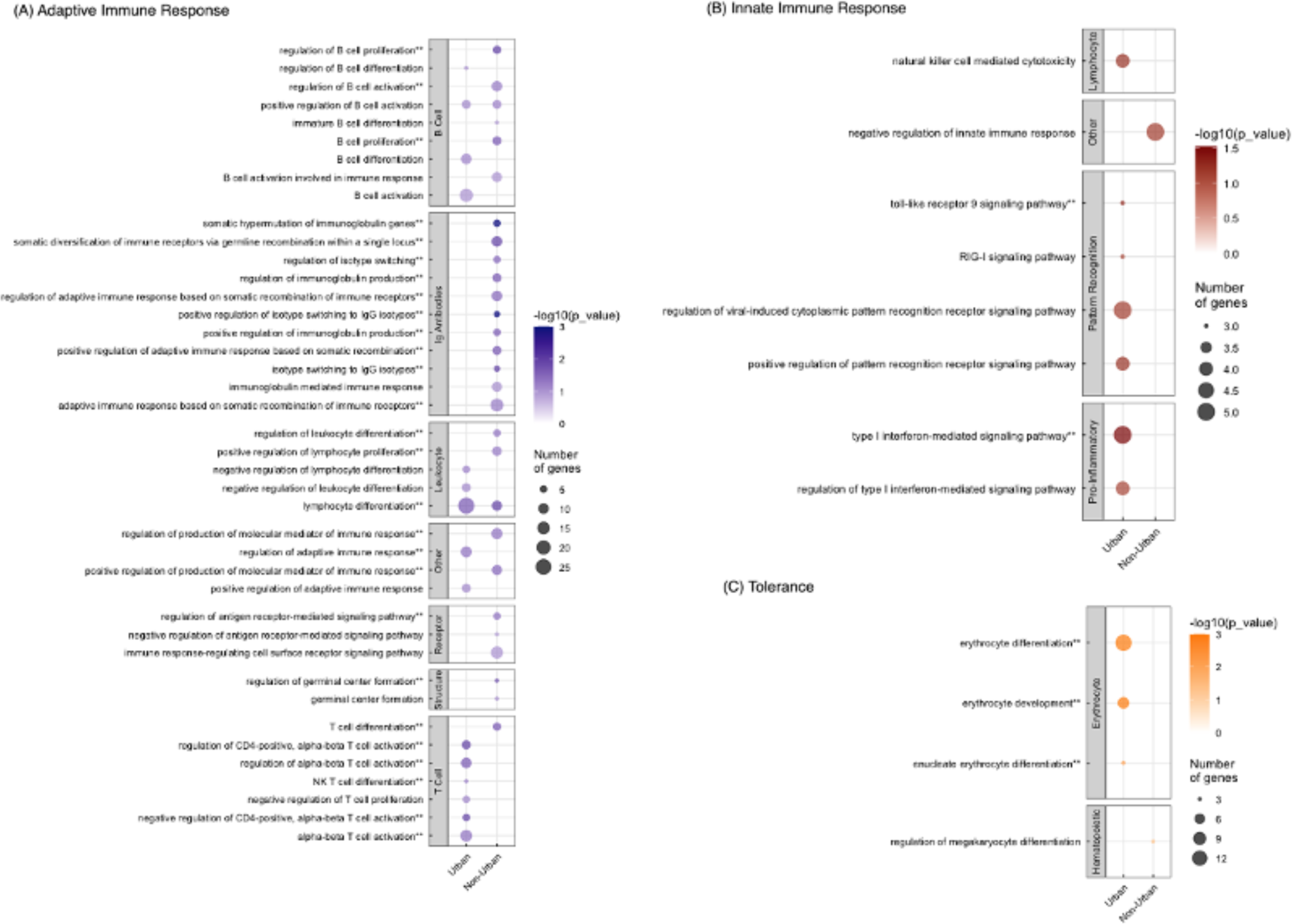
*Immune system process* Gene Ontology enrichment analysis of sham-fumigated urban (up-regulated) and non-urban (down-regulated) differentially expressed genes utilizing ClueGO. A) Adaptive Immune Response functional categories shown in purple were partitioned into seven sub-categories: B Cell, Ig Antibodies, Leukocyte, Receptor, Structure, T Cell and Other. B) Innate Immune Response categories, shown in red, were divided into four sub-categories: Lymphocyte, Pattern Recognition, Pro-Inflammatory and Other. C) Tolerance categories, shown in orange, were split into two sub-categories: Erythrocyte and Hematopoietic. Log10 (*P-*value) significance of unique and shared terms is depicted a color saturation gradient (** = *P-*value < 0.05), and number of genes supporting each ontology term is represented by circle size.

We next examined enrichment patterns related to *innate immunological resistance* (Fig. 4B). Urban sham-fumigated nestlings were enriched for both type I interferon mediated signaling (*P* < 0.03) and toll-like receptor 9 signaling (*P* < 0.05). Among the non-urban nestlings, only negative regulation of innate immune response (*P* < 0.07) was enriched.

Finally, we examined enrichment related to *tolerance* (Fig. 4C). Urban sham-fumigated nestlings had three significantly enriched terms specific to red blood cells, including erythrocyte differentiation (*P <* 0.001) and development (*P <* 0.001), as well as enucleate erythrocyte differentiation (*P* < 0.01). Among non-urban nestlings, regulation of megakaryocyte differentiation yielded a *P-*value less than 0.07.

### Co-expression Analysis

A differential co-expression analysis was conducted to provide novel insights on other traits that may explain gene expression patterns in nestling finches. Correlations in transcript levels across 12 traits (urban, sham-fumigated, hemoglobin level, glucose level, alive, δ^13^C, δ^15^N, C:N, days old, number of flies in nest, bill surface area and scaled mass) introduced 44 modules (File S3; Fig. S5-S8). Nestling survival (“alive” trait) presented the most significant correlations across the gene set, as illustrated by module A (*r* = 0.67, *P* < 0.0001) and module B (*r* = −0.82, *P* < 0.0001) respectively (Fig. 5). From the 2,338 total genes included in module A, 43 yielded 19 significantly enriched *immune system process* terms related to T-cell activation, lymphocyte differentiation, and negative regulation of hemopoiesis. Comparatively, 55 of the 3,422 genes in module B activated 21 immune pathways related to somatic hypermutation of immunoglobulin genes, leukocyte mediated cytotoxicity and interestingly, erythrocyte differentiation.

**Fig. 5.**
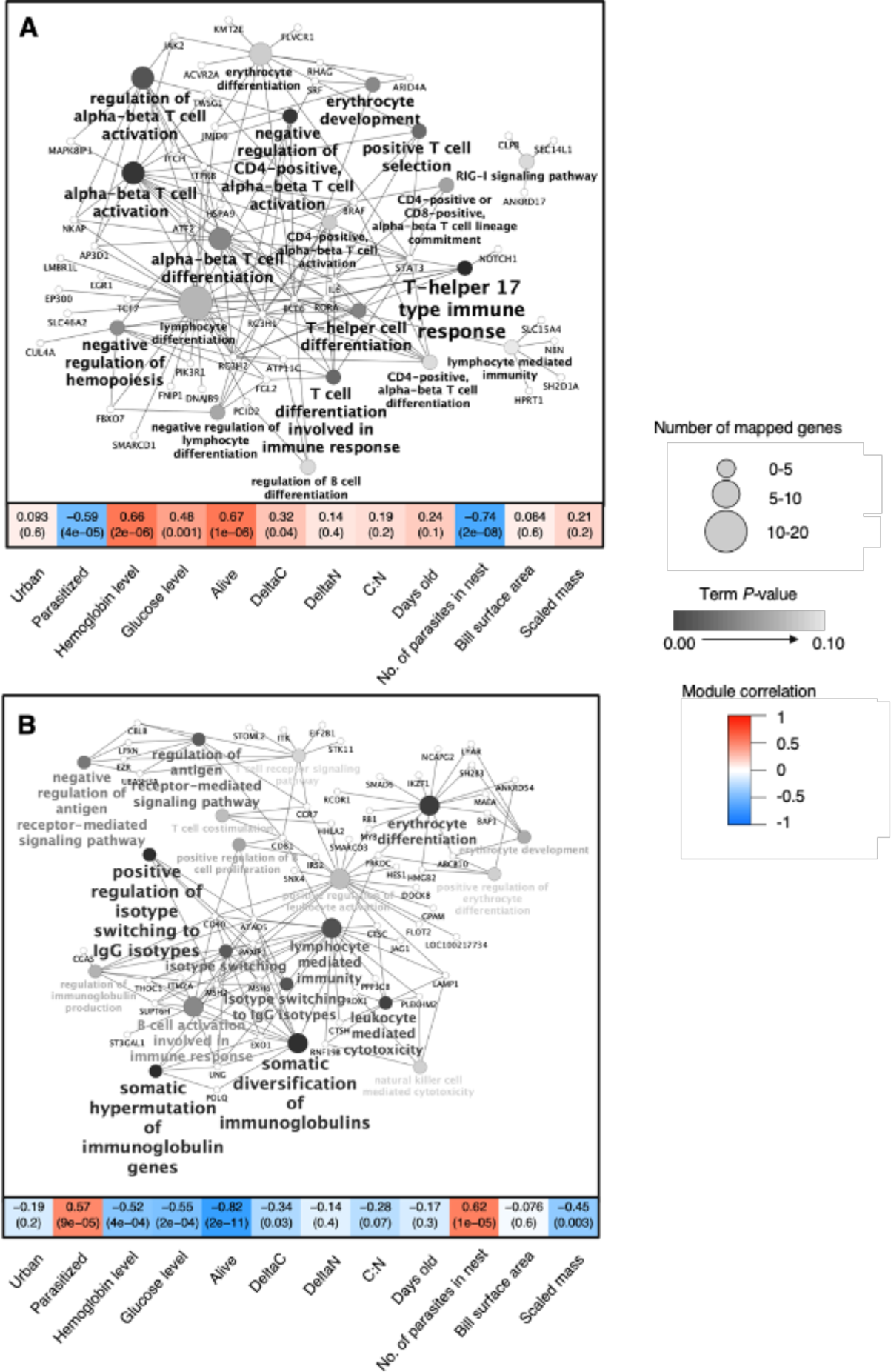
*Immune system process* Gene Ontology enrichment network of co-expression module A (top) and B (bottom). WGCNA trait correlation is depicted on a scale from 0 (white) to 1 (red), and negative trait correlation significance from 0 (white) to −1 (blue). WGCNA *P-*values are shown in parentheses. The co-expressed genes are represented with ClueGO in pathway form. The significance of enriched Gene Ontology terms is represented by the gray gradient. The size of the circle represents the number of genes associated with the enriched term.

Apart from mortality, other traits were strongly correlated with module A and B. Specifically, module A was negatively correlated with parasitism (*r* = −0.59, *P <* 0.0001) and nest parasite abundance (*r* = −0.74, *P* < 0.0001), and positively correlated with hemoglobin level (*r* = 0.66, *P* < 0.0001), glucose level (*r* = 0.48, *P* < 0.001) and δ^13^C (*r* = 0.32, *P <* 0.04). Specifically, module B was positively correlated with parasitism (*r* = 0.57, *P* < 0.0001) and nest parasite abundance (*r* = 0.62, *P* < 0.0001), and negatively correlated with hemoglobin level (*r* = −0.52, *P* < 0.0004), glucose level (*r* = −0.55, *P* < 0.0002), δ^13^C (*r* = −0.34, *P <* 0.03) and scaled mass (*r* = −0.45, *P* < 0.003). By comparison, although there were no notable correlations to urbanization in the described modules, the enriched immune pathways better explain patterns of expression observed in urban and non-urban sham-fumigated finches.

## Discussion

Our study found that urban living partially rescues nestling small ground finches from the lethal effects of parasitism by invasive avian vampire flies. Survival of sham-fumigated nestlings did not differ significantly between areas, which suggests that urbanization alone does not affect fledging success. Urban nests had significantly fewer flies than non-urban nests, such that even sham-fumigated urban nestlings survived. These results indicate that urban nestlings have effective resistance and tolerance mechanisms to deal with avian vampire flies. Differences in diet (measured using stable isotopes) between locations could explain why urban nestlings see fewer negative fitness consequences, but these results are correlative. Gene expression analyses revealed that resistance and tolerance mechanisms might underlie differences in parasite effects. Parasitized urban nestlings differentially expressed genes within pathways associated with tolerance and innate immunological resistance, compared to sham-fumigated non-urban nestlings. In contrast, sham-fumigated non-urban nestlings differentially expressed genes within pathways associated with adaptive immunological resistance. The gene expression results suggest that non-urban nestlings are investing in adaptive immunity, but that this type of response is not an effective defense mechanism against the parasite since few nestlings survived.

Overall, our field data suggest that the urban finch population is investing in resistance because urban nests had 40% fewer flies and higher nestling survival than non-urban nests. Gene expression profiles showed that urban nestlings differentially express innate immune genes associated with pro-inflammatory cytokines, specifically type-1 interferons (IFNs), compared to non-urban nestlings. When larval flies chew through the skin of their hosts, effective inflammation by the host (thickening the skin and restricting blood flow) can prevent ectoparasite feeding (reviewed in Owens et al. 2010). Although other avian urbanization studies have not observed changes in IFNs, they have found other enhanced innate immune responses within urban populations. For example, Watson et al. (2017) observed overrepresented genes involved in the secretion and receptor-binding of cytokines in urban great tits (*Parus major*) compared to non-urban tits. Additionally, urban nestling black sparrowhawks (*Accipiter melanoleucus*) had a stronger innate response to an immune challenge compared to non-urban nestlings (Nwaogu et al. 2023). These studies provide evidence that urban living could confer an advantage against parasitism. Our study further supports this idea, as shown by the heightened innate immune response in urban nestlings, and also links this response with a decrease in parasite abundance.

Most studies suggest that IFNs are largely involved in resistance to viruses (Fensterl & Sen, 2009; Katze et al. 2002). An experimental study (e.g., with avian vampire fly-specific vaccination challenges) is still needed to determine whether IFNs are specifically involved in the inflammatory response to avian vampire flies because it is possible that finches are actually responding to another parasite or pathogen (e.g., virus). For example, adult finches in Puerto Baquerizo Moreno, San Cristóbal Island, are susceptible to infection by the invasive avian pox virus (Lynton-Jenkins et al. 2021). A recent study suggests that pox-infected adult finches upregulate expression of interferon pathways (McNew et al. 2022), which could explain the expression of IFN seen in urban sham-fumigated nestlings. One interesting possibility is that infection by pox could also be conferring resistance to avian vampire flies. This potential explanation requires further study, but could provide insight into the disease dynamics of co-infecting invasive parasites of Galápagos birds (Wikelski et al. 2004).

Although sham-fumigated non-urban nestlings did not exhibit significantly enriched innate immune pathways, they did differentially express genes involved in pathways related to the adaptive immune response. Specifically, T-cell, B-cell, and Ig pathways were expressed, but without successfully conferring resistance since almost all of the nestlings died. This upregulation could be the final effort by the finches’ immune system to deal with the parasite before it becomes physiologically costly. In contrast, urban finch nestlings expressed more adaptive immune pathways related to lymphocyte and T-cell differentiation, compared to non-urban nestlings. One explanation for why urban nestlings had higher survival compared to non-urban nestlings is that resistance is heightened when the innate and adaptive immune system are activated simultaneously (Palm & Medzhitov 2007).

Regardless of the mechanism, a central question is why urban finches are more resistant to avian vampire flies than non-urban finches. Immunological resistance can be conditionally dependent and only hosts in good condition may be able to resist parasites (Cornet, Bichet, Larcombe, Faivre, & Sorci, 2014; Howick & Lazzaro, 2014; Lochmiller & Deerenberg, 2000; Sheldon & Verhulst, 1996; Sternberg et al., 2012; Svensson, Råberg, Koch, & Hasselquist, 1998). In a native host-parasite system, eastern bluebirds (*Sialia sialis*) are both tolerant and resistant to nest flies and the investment in defenses depends on whether supplemental food is available (resistance) or not (tolerance; Knutie 2020). Urban areas in the Galápagos provide a reliable source of human food for animals, including finches (De León et al. 2018). For example, some restaurants have outdoor dining, where food falls off of tables and finches feed on the tables (Fig. S9). Our stable isotope results corroborate the idea that diet differs between urban and non-urban nestlings. In fact, higher δ^15^N values suggest a diet rich in protein, which could include food items such as chicken and fish, found at outdoor restaurants in town. Higher δ^15^N values also correlated with lower parasite intensities, which supports the idea that diet might influence host defenses. For example, meat has higher protein concentrations than plants and supplemented protein can increase the concentration of cellular immune cells (e.g., eosinophils, globule leukocytes and mast cells) (reviewed in Coop and Kyriazakis, 2001). Although our results suggest that urban finches are more physiologically resistant to the flies than non-urban finches, another potential explanation is that nest material affects the survival of flies. For example, urban Darwin’s finches incorporate cigarette butts into their nests (Harvey et al. 2021), which can affect parasite abundance in other systems (Suarez-Rodriguez et al. 2013). Future studies could explore the idea that nest material could contribute to increased resistance in urban finches.

Some urban nests had 100% nestling survival despite high parasite abundances (up to 48 flies), indicating that these urban nestlings are tolerant of flies. Studies have demonstrated that low finch survival in response to vampire flies is likely related to exsanguination (i.e., high blood loss) (Fessl et al. 2006). One explanation for increased tolerance is that urban nestlings have effective blood recovery when sham-fumigated and thus tolerate parasitism. This hypothesis is corroborated with our results that urban finches have more hemoglobin (oxygenated blood) when sham-fumigated compared to non-urban birds. Gene expression profiles suggest that urban nestlings differentially expressed genes within pathways associated with red blood cell production compared to non-urban nestlings when sham-fumigated. Furthermore, nestlings that survived had higher gene expression of blood cell production than nestlings that died, which was observed across urban and non-urban sites. Galápagos mockingbirds are also relatively tolerant of avian vampire flies (Knutie et al. 2016), but this tolerance is lost during dry years with low food availability (McNew et al. 2019). The working hypothesis is that mockingbirds are a larger-bodied species and that larger hosts might be more tolerant of avian vampire flies (McNew and Clayton 2018). However, our study suggests that even smaller-bodied hosts can tolerate avian vampire flies, and there is likely an alternative explanation. The hormone erythropoietin is produced primarily by the kidneys and works together with iron to induce erythropoiesis, which is the production of red blood cells. One explanation for the lack of tolerance in non-urban nestlings is that these nestlings are not receiving sufficient amounts of iron or are not producing enough erythropoietin. Urban nestlings have higher δ^15^N values, which is likely because they are feeding on more iron-rich meat products than non-urban birds. Thus, this difference in diet could explain the increased tolerance related to red blood cell recovery, but requires further studies on iron and nestling endocrinology. Finally, although non-urban nestlings did not exhibit any significantly enriched pathways related to tolerance, we found that adenosine deaminase (ADA) was significantly up-regulated. Interestingly, overexpression of ADA in red blood cells has been shown to cause hemolytic anemia (Chen & Mitchell 1994), which could be additionally responsible for non-urban finch mortality.

Although survival of urban nestlings in response to flies is higher than non-urban nestlings, some urban nests still failed. Therefore, not all urban finches have effective defenses against avian vampire flies. Our urban field location is environmentally heterogeneous with nests found in areas of high tourist activity, residential areas, a naval base, and an airport. Our spatial analysis did not find any distinct patterns for parasite abundance or fledging success related to these landmarks. However, human food type, abundance, and reliability vary across these locations and over time. Future research should consider how the heterogeneity in urban areas may favor more phenotypic plasticity in effective defenses and/or hinder adaptations in the population.

Heterogeneity across different urban areas could also impact finch-fly interactions on other islands in the Galapagos. Due to logistical limitations, replication of sites was not possible across islands. However, site replication would be helpful to understand whether context dependency exists across different urban areas. The Galapagos Islands have two other major towns, including Puerto Ayora of Santa Cruz Island with approximately 12,000 human residents and Puerto Villamil of Isabela Island with approximately 2,000 human residents (Memoria Estadística Galápagos 2017). Human population size often scales with anthropogenic features (e.g., artificial light at night, density of restaurants, number of vehicles) in urban areas, which can proportionally affect bird species (reviewed in Isaksson 2018). Thus, replicating our study in towns on other islands in the Galapagos could provide insight into whether the scale of urbanization impacts finch-fly interactions differently.

Naïve hosts are thought to lack defenses against novel parasites, which can cause species declines and extinctions (Daszak et al. 2000, Keesing et al. 2010, Atkinson and Lapointe 2009). Over the past decade, studies have found that invasive avian vampire flies can cause up to 100% mortality in Darwin’s finch species across islands in the Galápagos (O’Connor et al. 2010, Koop et al. 2010, Koop et al. 2013, O’Connor et al. 2014; Klendorfer and Dudaniec 2016; Knutie et al. 2016, McNew and Clayton 2018, Addesso et al. 2021). If humans cannot effectively control the fly (e.g., Knutie et al. 2014) or host species do not evolve or develop defenses, some species may even go extinct (Fessl et al. 2010, Koop et al. 2013b). However, our study provides experimental evidence that an urban population of finches is relatively well-defended against avian vampire flies. Indeed, additional studies of urban and non-urban populations across years are needed to determine the magnitude of the urban effect, but preliminary data from other years (2022, 2023) and populations (Puerto Chino, San Cristobal) suggests that this pattern is observed beyond our study (SAK prelim. data). To be clear, we are not suggesting that the Galápagos Islands should be urbanized to increase defense against flies. Instead, the goal of our study is to demonstrate that some finches can defend themselves against the flies through various mechanisms, which can provide insight into management strategies (Ohmer et al. 2021).

## Acknowledgements

We would like to thank the Galápagos Science Center and the Galápagos National Park for support. Specifically, we thank Karla Vasco from the Galápagos Science Center for her assistance and logistical support in the laboratory. We thank Bo Reese for assistance at the UConn Center for Genomic Innovation and Corrine Arthur for help with field work. The work was supported by start-up funds and an Institute for Systems Genomics grant from the University of Connecticut, a National Science Foundation (NSF) Grant (DEB-1949858), and a National Geographic Grant to SAK, an NSF grant to JW (DBI-1943371), an Explorers Club Mamont Scholar Grant, an Animal Behavior Society Student Research Grant, a Whetten Travel award from El Instituto at the University of Connecticut, and a University of Connecticut Department of Ecology and Evolutionary Biology Zoology 2019 Award to GJV. All bird handling and work was conducted according to approved University of Connecticut IACUC (Institutional Animal Care and Use Committee) protocols (No. A17-044). Our work was done under GNP permits PC 28-19 and Genetic Access permit MAE-DNB-CM-2016-0041. Authors declare that they have no competing interests.

## Data Availability Statement

Supplemental files 1-3 and details of full analysis, including intermediate files and supporting scripts, are publicly available at: https://gitlab.com/PlantGenomicsLab/galapagos-finch-rna-seq. All raw data are available on FigShare (DOI: available upon acceptance) and sequences have been uploaded to GenBank (BioProject accession number: PRJNA930453).

## Author contributions

Conceptualization: SAK; Methodology: SAK, CW, GJV, JLW; Analyses: SAK, CW, GJV, ML, JLW; Investigation: SAK, CW, GJV, LA, CA, TBV, AS, JAH, JLW; Visualization: SAK, CW, ML, JLW; Funding acquisition: SAK, GJV, JLW; Project administration: JC, SAK; Supervision: SAK, JLW; Writing – original draft: SAK, CW, GJV, ML, JLW; Writing – review & editing: All authors.

