## Supplemental material for "Urban living can rescue Darwin’s finches from the lethal effects of invasive vampire flies"

Supplementary Materials for Knutie, Webster et al.

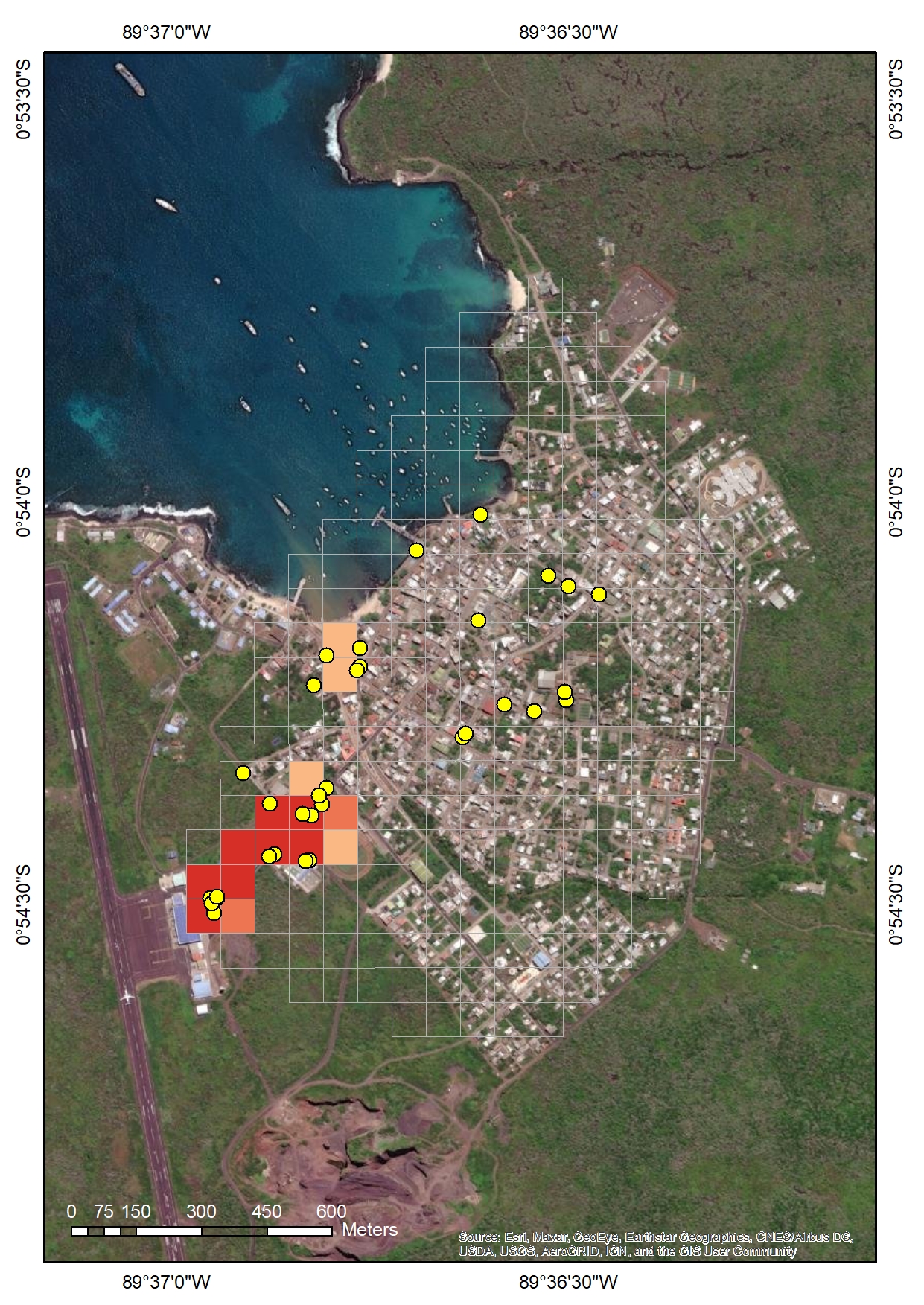

Fig. S1. Results of optimized hot-spot analysis (conducted in ArcGIS 10.8) for urban nests of finches in San Cristóbal based only on the spatial patterns of the nest locations. Yellow dots represent nest locations (*n* = 37) and red grid cells represent >90% confidence of the cell being a hotspot for nest locations, which are situated toward the southwest part of the city near the airport.

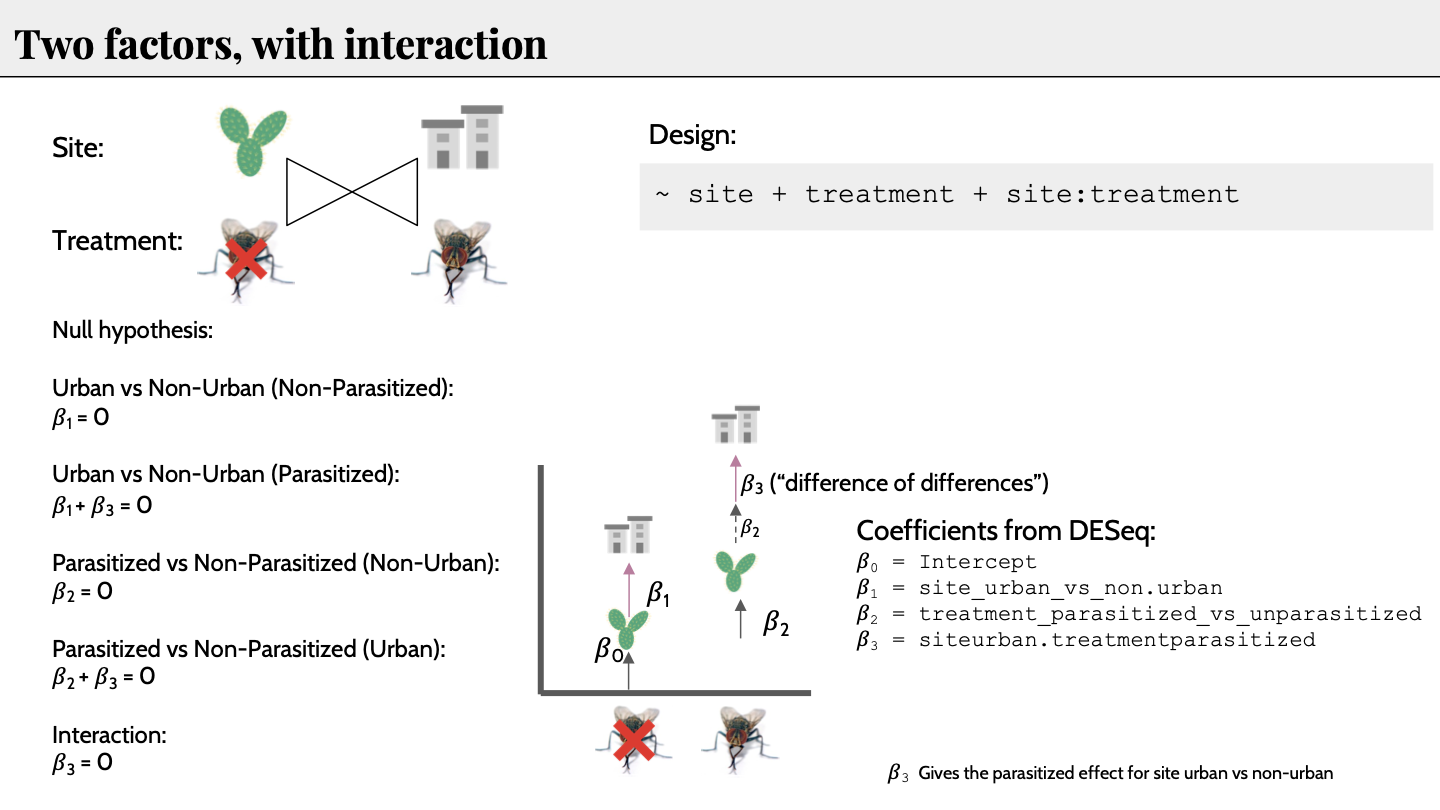

Figure S2. DESeq2 pairwise-differential expression with two factors and interaction design. Urban (plant) and non-urban location (building) compared to sham-fumigated (fly) and fumigated (exed fly). Coefficients from DESeq2, 𝛽0, 𝛽1, 𝛽2, and 𝛽3 used to define design and expected null hypotheses for each pairwise comparison.

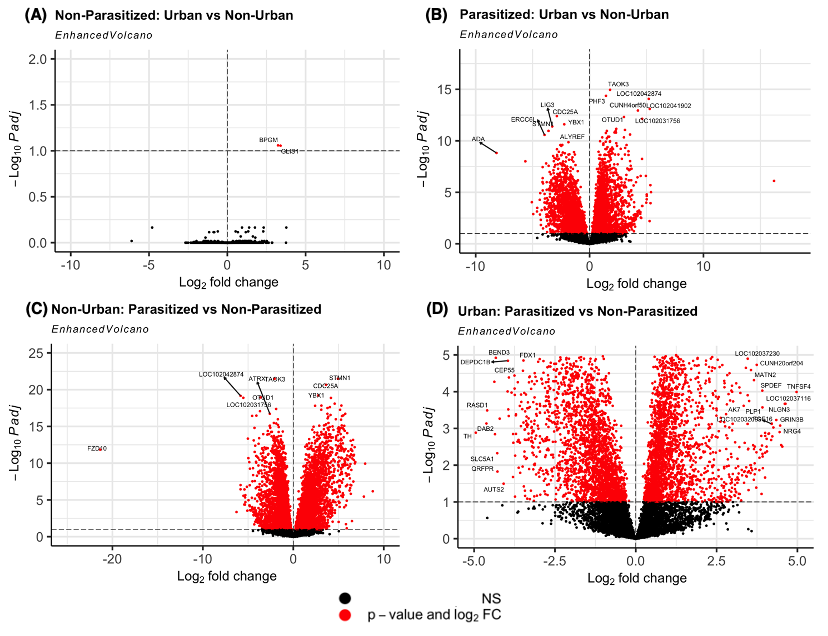

Figure S3. Volcano plot of differentially expressed genes from each pairwise comparison: (A) Sham-Fumigated: Urban vs Non-Urban, (B) Fumigated: Urban vs Non-Urban, (C) Urban: Sham-Fumigated vs Fumigated, and (D) Non-Urban: Sham-Fumigated vs Fumigated. The red points show differentially expressed genes of high statistical significance (*-*Log10 *P-*adj > 1.0, y-axis) and fold change (FC > 0, x-axis). The black points show genes that are not statistically significant (*-*Log10 *P-*adj < 1.0).

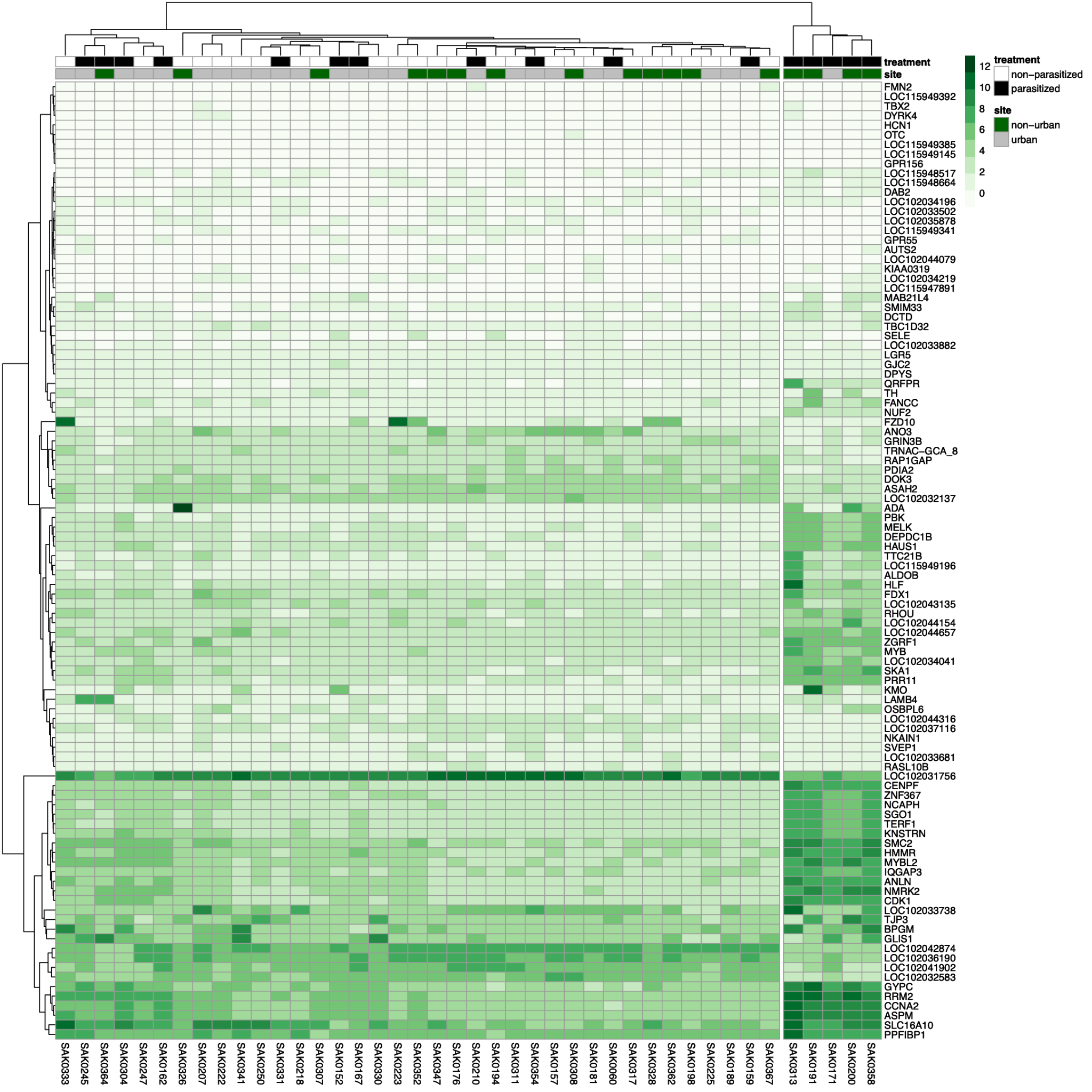

Figure S4: Heatmap of top 100 differentially expressed genes across 42 nestling samples that are either sham-fumigated (black) or fumigated (white), and urban (gray) or non-urban (dark green). Significance of gene expression per each individual depicted by white (low) to green (high) color gradient scaling from 0 to 12. Samples that exhibit similar patterns of expression are clustered together.

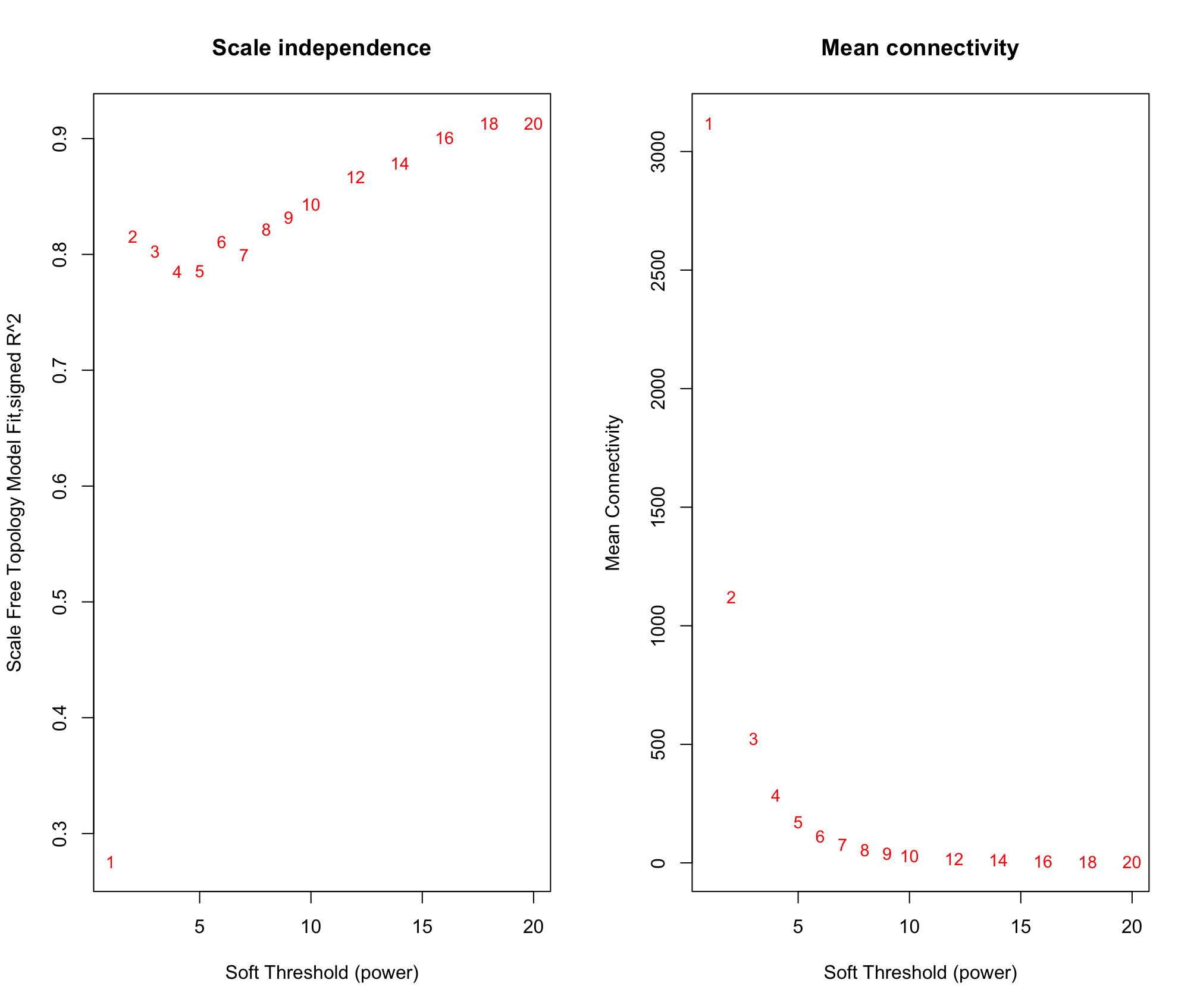

Figure S5. Network topology of different soft-thresholding powers. The left panel shows scale free topology (y-axis) as a function of soft thresholding power (x-axis). The right panel depicts mean connectivity (x-axis) as a function of soft-thresholding power (x-axis). The soft thresholding power 6 was chosen to construct the gene network and identify modules in WGCNA.

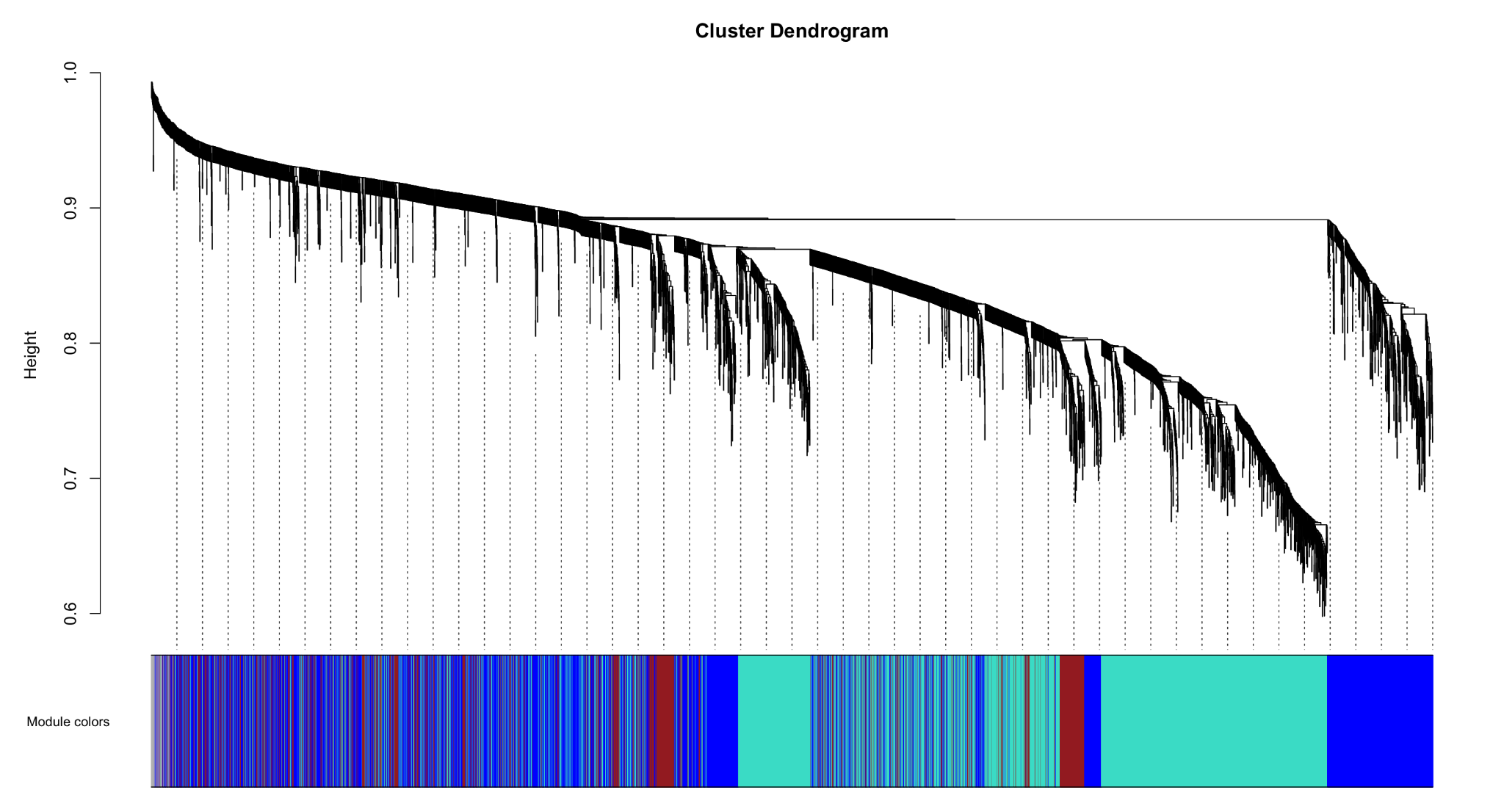

Figure S6. Cluster dendrogram of nestling genes. Gene tree mapped to co-expression modules (colors) produced by hierarchical clustering of genes by topological overlaps in RNA-expression. X-axis shows all genes in the gene set, and y-axis shows the height of the gene tree.

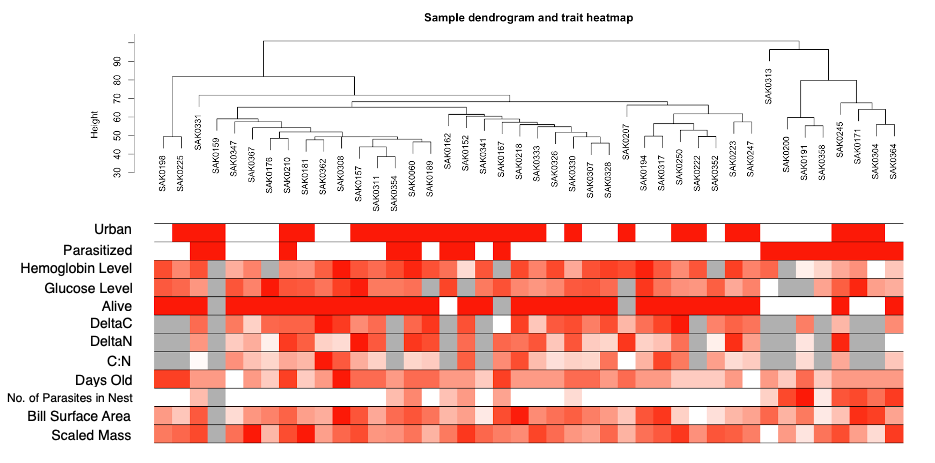

Figure S7. Sample dendrogram and trait heatmap. Nestling samples displaying similar expression patterns were clustered together on the tree. Height (y-axis) was used to detect outliers. SAK0313 could be identified as a possible outlier, however we chose to keep all samples in order to detect correlations in morality. Associated binary (solid) and continuous (gradient) traits are depicted under each sample (red: present, white: absent, gray: NA).

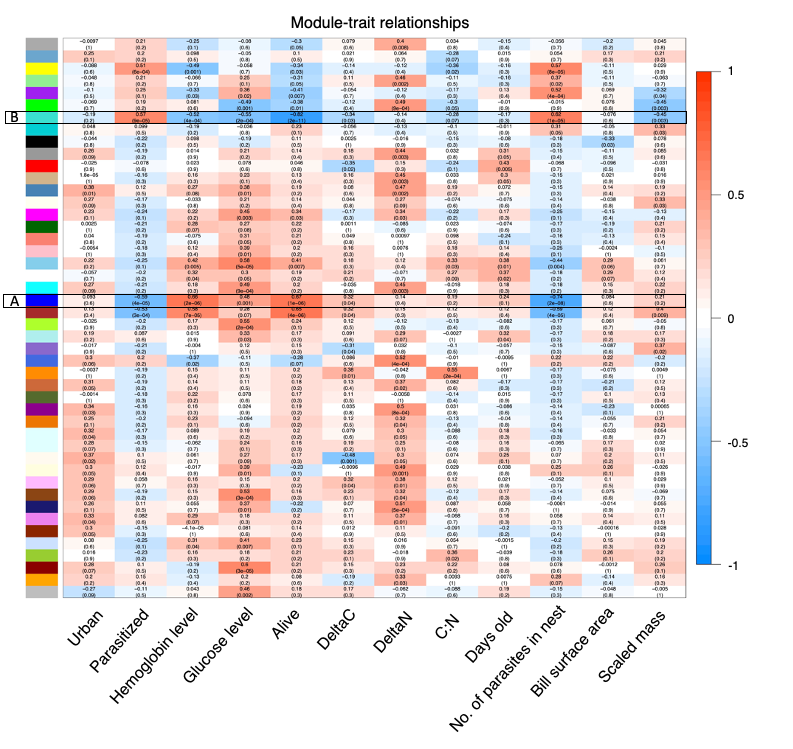

Figure S8. Co-expression module trait relationships. Genes binned into color modules (left) by nestling expression patterns. Pearson correlation of each module is depicted by a blue (negative) and red (positive) color gradient, and *P-*value in parentheses. Module A and B referenced in the manuscript are shown in a black box.

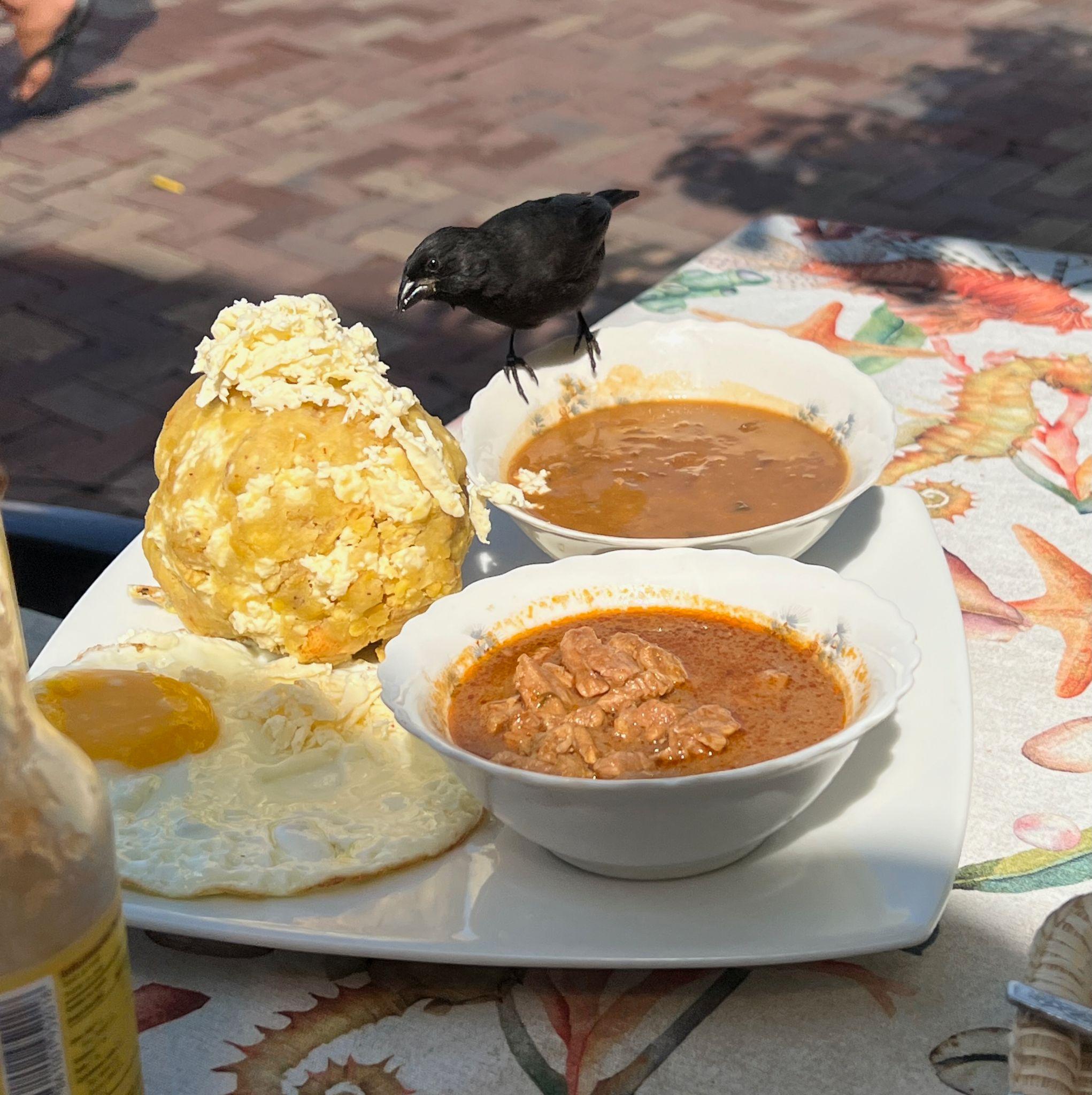

Figure S9. Male small ground finch foraging on a restaurant table in Puerto Baquerizo Moreno, San Cristobal. Food on the plate include cheese, bolon (plantains and cheese), beef, beans, and an egg. Photo by S. Knutie.

Table S1. Individual data for nestling morphometrics and physiology across locations (NU = Non-urban, U = Urban) and treatments (F = Fumigated, SF = Sham-fumigated). Nest parasite abundance is abbreviated “Nest PA”.

| Nest ID | Band number | Location | Treatment | Days Old | Julian hatch day | Nest PA | Bill length (mm) | Bill depth (mm) | Bill width (mm) | Tarsus length (mm) | Primary length (mm) | Mass (g) | Hemoglobin (g/dL) | Glucose (mg/dL) | Scaled mass index | Bill surface area |
| --- | --- | --- | --- | --- | --- | --- | --- | --- | --- | --- | --- | --- | --- | --- | --- | --- |
| 71 | SAK0351 | NU | F | 4 | 118 | 0 | 3.35 | 4.1 | 5 | 13.3 | 5.3 | 7.1 | NA | NA | 9.97 | 23.94 |
| 38b | SAK0249 | U | F | 4 | 124 | 0 | 3.7 | 4.3 | 4 | 12.3 | 4.1 | 5.3 | NA | NA | 8.56 | 24.12 |
| 3 | SAK0175 | NU | F | 5 | 65 | 0 | 4.1 | 5.1 | 5 | 14.4 | 6.35 | 7.1 | NA | NA | 8.64 | 32.52 |
| 65 | SAK0347 | NU | F | 5 | 118 | 0 | 3.95 | 3.9 | 4.8 | 14.6 | 8.1 | 7.4 | 6.8 | 286 | 8.79 | 26.99 |
| 38b | SAK0247 | U | F | 5 | 124 | 0 | 4.4 | 4.1 | 5.5 | 14.6 | 5.25 | 7.1 | NA | 223 | 8.43 | 33.18 |
| 38b | SAK0248 | U | F | 5 | 124 | 0 | 4.2 | 3.6 | 5.2 | 13.7 | 3.6 | 5.9 | 8.2 | 229 | 7.86 | 29.03 |
| 38b | SAK0246 | U | F | 5 | 124 | 0 | 4.1 | 4.25 | 5.2 | 13.9 | 5.85 | 6.1 | NA | NA | 7.91 | 30.43 |
| 44c | SAK0251 | U | SF | 5 | 122 | 48 | 3.2 | 3.05 | 4.4 | 9.15 | 1.25 | 3.4 | NA | NA | 9.34 | 18.72 |
| 44c | SAK0252 | U | SF | 5 | 122 | 48 | 3.5 | 2.9 | 4.45 | 9.25 | 0.8 | 3.2 | NA | NA | 8.62 | 20.20 |
| 44c | SAK0253 | U | SF | 5 | 122 | 48 | 3.4 | 3.5 | 4.4 | 10.55 | 1.85 | 4.6 | NA | NA | 9.79 | 21.10 |
| 3 | SAK0177 | NU | F | 6 | 65 | 0 | 4.15 | 4.5 | 5.4 | 15.3 | 6.4 | 8.7 | NA | 347 | 9.50 | 32.27 |
| 3 | SAK0176 | NU | F | 6 | 65 | 0 | 4.75 | 4.9 | 5.1 | 17.45 | 9.7 | 8.4 | NA | 379 | 7.24 | 37.31 |
| 27 | SAK0195 | NU | F | 6 | 73 | 0 | 5 | 4.5 | 5.5 | 16.55 | 9 | 9.4 | NA | 332 | 8.91 | 39.27 |
| 27 | SAK0196 | NU | F | 6 | 73 | 0 | 5.35 | 4.85 | 5.8 | 17.45 | 9.6 | 10.2 | 11.6 | NA | 8.75 | 44.75 |
| 33 | SAK0200 | NU | SF | 6 | 76 | 41 | 4.5 | 4.3 | 4.7 | 16.2 | 11.05 | 8 | NA | NA | 7.88 | 31.81 |
| 34 | SAK0316 | NU | F | 6 | 93 | 0 | 4.85 | 4.75 | 5.9 | 17.8 | 10.75 | 9.8 | NA | 242 | 8.15 | 40.57 |
| 34 | SAK0318 | NU | F | 6 | 93 | 0 | 4.65 | 5.05 | 6.3 | 16.2 | 9.5 | 9.5 | NA | NA | 9.36 | 41.45 |
| 65 | SAK0348 | NU | F | 6 | 118 | 0 | 4.8 | 4.25 | 5.1 | 16.5 | 10.55 | 9.5 | NA | 324 | 9.06 | 35.25 |
| 65 | SAK0349 | NU | F | 6 | 118 | 0 | 3.2 | 2.85 | 3.6 | 14.8 | 9.3 | 8.8 | NA | NA | 10.20 | 16.21 |
| 71 | SAK0350 | NU | F | 6 | 118 | 0 | 3.5 | 4.1 | 5.15 | 15.2 | 7.6 | 8.6 | NA | 241 | 9.50 | 25.43 |
| 71 | SAK0352 | NU | F | 6 | 118 | 0 | 3.7 | 4.05 | 4.95 | 14.9 | 7.3 | 8.2 | NA | 277 | 9.39 | 26.15 |
| 72 | SAK0358 | NU | SF | 6 | 120 | 5 | 3.9 | 3.7 | 4.5 | 12.6 | 5.25 | 5.6 | 5.6 | 185 | 8.67 | 25.12 |
| 12 | SAK0180 | U | F | 6 | 67 | 0 | 4.2 | 4.4 | 5.5 | 15.55 | 8.9 | 9.1 | 8.2 | 250 | 9.65 | 32.66 |
| 12 | SAK0181 | U | F | 6 | 67 | 0 | 4.3 | 4.7 | 5.3 | 15.8 | 10 | 10.4 | 7.7 | 280 | 10.72 | 33.77 |
| 12 | SAK0179 | U | F | 6 | 67 | 0 | 3.7 | 4.5 | 4.6 | 11.6 | 4.65 | 6.7 | NA | NA | 12.03 | 26.44 |
| 16b | SAK0224 | U | F | 6 | 96 | 0 | 3.9 | 4.05 | 4.95 | 15.7 | 8.75 | 9.2 | NA | NA | 9.59 | 27.57 |
| 24 | SAK0173 | U | F | 6 | 65 | 0 | 3.5 | 4.2 | 4.5 | 13.3 | 5.75 | 6.4 | NA | NA | 8.99 | 23.92 |
| 24 | SAK0174 | U | F | 6 | 65 | 0 | 4.1 | 3.5 | 4.6 | 15.15 | 7 | 8 | NA | NA | 8.89 | 26.08 |
| 24b | SAK0341 | U | F | 6 | 114 | 0 | 3.55 | 4.3 | 4.8 | 11.5 | 3.8 | 4.9 | 9.8 | 241 | 8.93 | 25.37 |
| 35c | SAK0250 | U | F | 6 | 123 | 0 | 3.75 | 4.6 | 5 | 14.85 | 8.2 | 8.7 | 7.9 | 208 | 10.02 | 28.27 |
| 38a | SAK0215 | U | F | 6 | 88 | 0 | 4.6 | 4.75 | 5.85 | 14 | 7.05 | 6.5 | NA | NA | 8.32 | 38.30 |
| 44b | SAK0220 | U | F | 6 | 96 | 0 | 3.3 | 2.9 | 4.8 | 14 | 7.1 | 6.8 | 8.5 | 177 | 8.71 | 19.96 |
| 44b | SAK0222 | U | F | 6 | 96 | 0 | 3.3 | 3.7 | 5 | 14.5 | 5.8 | 6.7 | 9.7 | 199 | 8.06 | 22.55 |
| 44b | SAK0221 | U | F | 6 | 96 | 0 | 3.6 | 3.5 | 4.6 | 12.9 | 3.95 | 6.3 | NA | NA | 9.35 | 22.90 |
| 53b | SAK0243 | U | SF | 6 | 121 | 38 | 4.2 | 3.75 | 4.7 | 14 | 5.3 | 7.8 | NA | NA | 9.99 | 27.87 |
| 03b | SAK0357 | NU | SF | 7 | 119 | 22 | 4.5 | 3.7 | 4.85 | 14.05 | 5.3 | 6.8 | NA | 247 | 8.65 | 30.22 |
| 03b | SAK0356 | NU | SF | 7 | 119 | 22 | 4.25 | 4.3 | 4.75 | 13.15 | 5 | 6 | 6.7 | NA | 8.60 | 30.21 |
| 11 | SAK0193 | NU | SF | 7 | 72 | 53 | 4.65 | 4.4 | 5.25 | 16.5 | 9.1 | 7.9 | NA | NA | 7.53 | 35.24 |
| 27 | SAK0194 | NU | F | 7 | 73 | 0 | 5.3 | 4.8 | 5.65 | 17.65 | 11 | 10.4 | NA | NA | 8.79 | 43.50 |
| 33 | SAK0301 | NU | SF | 7 | 76 | 41 | 4.5 | 3.8 | 4.6 | 16.2 | 10.85 | 9 | 4.7 | NA | 8.87 | 29.69 |
| 34 | SAK0317 | NU | F | 7 | 93 | 0 | 5.4 | 5.25 | 5.95 | 17.65 | 13.05 | 10.9 | 9.6 | NA | 9.21 | 47.50 |
| 46 | SAK0327 | NU | F | 7 | 105 | 0 | 5.15 | 4.8 | 5.8 | 15.75 | 8.7 | 8.75 | NA | 218 | 9.07 | 42.87 |
| 46 | SAK0326 | NU | F | 7 | 105 | 0 | 4.8 | 5.25 | 5.3 | 16.45 | 11 | 9.25 | 10.3 | NA | 8.87 | 39.77 |
| 47 | SAK0313 | NU | SF | 7 | 85 | 0 | 5.1 | 4.7 | 5.5 | 14.5 | 8.7 | 4.8 | NA | 67 | 5.77 | 40.86 |
| 50 | SAK0368 | NU | F | 7 | 123 | 0 | 4.5 | 5.3 | 5.5 | 16.4 | 9.3 | 11.1 | NA | 228 | 10.70 | 38.17 |
| 50 | SAK0367 | NU | F | 7 | 123 | 0 | 4.1 | 5 | 5.9 | 16.8 | 10.02 | 11.7 | 8.2 | NA | 10.80 | 35.10 |
| 54 | SAK0328 | NU | F | 7 | 105 | 0 | 4.7 | 4.65 | 4.65 | 16.1 | 11 | 9 | 10.5 | 255 | 8.97 | 34.33 |
| 59 | SAK0364 | NU | SF | 7 | 123 | 44 | 4.3 | 5 | 4.8 | 13.85 | 8.05 | 7.6 | NA | 173 | 9.92 | 33.10 |
| 59 | SAK0366 | NU | SF | 7 | 123 | 44 | 4.4 | 5 | 4.55 | 15.4 | 9.2 | 7.4 | NA | NA | 7.99 | 33.00 |
| 59 | SAK0365 | NU | SF | 7 | 123 | 44 | 4.4 | 4.6 | 5.1 | 14.5 | 10.09 | 6.5 | 6 | 223 | 7.82 | 33.52 |
| 73 | SAK0362 | NU | F | 7 | 121 | 0 | 4.3 | 4.8 | 4.7 | 15.8 | 10.05 | 7.5 | 9.4 | 243 | 7.73 | 32.08 |
| 1 | SAK0162 | U | SF | 7 | 51 | 13 | 4.45 | 3.45 | 5.5 | 17 | 13.2 | 9.5 | NA | NA | 8.59 | 31.28 |
| 5 | SAK0156 | U | F | 7 | 47 | 0 | 4.7 | 4.4 | 5.9 | 17.1 | 13.5 | 9.8 | 9.8 | 216 | 8.76 | 38.02 |
| 5 | SAK0158 | U | F | 7 | 47 | 0 | 4.45 | 4.25 | 5.7 | 16.3 | 10.7 | 8.3 | NA | NA | 8.09 | 34.78 |
| 05c | SAK0355 | U | SF | 7 | 118 | 14 | 4.9 | 4.65 | 6.1 | 17 | 11.6 | 8.4 | 8.8 | 226 | 7.59 | 41.37 |
| 05c | SAK0353 | U | SF | 7 | 118 | 14 | 4.6 | 4.6 | 5.4 | 15.6 | 10.7 | 8.4 | NA | 239 | 8.86 | 36.13 |
| 05c | SAK0354 | U | SF | 7 | 118 | 14 | 5 | 4.6 | 5.6 | 17 | 11.95 | 8.9 | 10 | NA | 8.04 | 40.06 |
| 8 | SAK0152 | U | SF | 7 | 44 | 22 | 4.4 | 4.3 | 5.5 | 15.7 | 10 | 9.6 | NA | NA | 10.01 | 33.87 |
| 8 | SAK0155 | U | SF | 7 | 44 | 22 | 5 | 4.3 | 5.9 | 16.6 | 13.3 | 10.8 | NA | NA | 10.19 | 40.06 |
| 9 | SAK0190 | U | F | 7 | 70 | 0 | 4.7 | 4.3 | 5.6 | 17.9 | 11.5 | 8.2 | NA | NA | 6.76 | 36.54 |
| 16 | SAK0172 | U | SF | 7 | 64 | 41 | 5.5 | 5.35 | 6.65 | 16.9 | 13.2 | 7.9 | 7.7 | 347 | 7.22 | 51.84 |
| 16 | SAK0171 | U | SF | 7 | 64 | 41 | 5.2 | 5.35 | 6.55 | 16.7 | 13.15 | 8.3 | NA | 356 | 7.74 | 48.60 |
| 16b | SAK0223 | U | F | 7 | 96 | 0 | 4.2 | 4.3 | 5.05 | 16.75 | 10.03 | 9.8 | 10.6 | 179 | 9.10 | 30.84 |
| 20 | SAK0166 | U | SF | 7 | 62 | 10 | 4.6 | 3.6 | 5.25 | 15.8 | 10.6 | 8.8 | NA | NA | 9.07 | 31.97 |
| 26 | SAK0303 | U | SF | 7 | 77 | 31 | 4.7 | 4.8 | 5.9 | 15.9 | 8.3 | 5.8 | NA | 162 | 5.91 | 39.50 |
| 26 | SAK0305 | U | SF | 7 | 77 | 31 | 4.4 | 4.8 | 5.65 | 14.8 | 7.65 | 4.8 | 4.3 | 186 | 5.56 | 36.11 |
| 26 | SAK0304 | U | SF | 7 | 77 | 31 | 5.4 | 5 | 5.9 | 16.7 | 9 | 7 | NA | NA | 6.53 | 46.23 |
| 38a | SAK0218 | U | F | 7 | 88 | 0 | 5.4 | 5.4 | 6.6 | 17.8 | 12.2 | 9.75 | 10.3 | 221 | 8.11 | 50.89 |
| 38a | SAK0216 | U | F | 7 | 88 | 0 | 5.85 | 5 | 5.8 | 17.9 | 12.1 | 9.05 | NA | NA | 7.46 | 49.62 |
| 49 | SAK0331 | U | SF | 7 | 106 | 14 | 5.15 | 5 | 5.5 | 16.35 | 9.7 | 10 | 9.9 | 204 | 9.69 | 42.47 |
| 49 | SAK0332 | U | SF | 7 | 106 | 14 | 4.4 | 5.05 | 5.55 | 14.9 | 8.3 | 8.2 | NA | NA | 9.39 | 36.63 |
| 52 | SAK0333 | U | F | 7 | 106 | 0 | 4.85 | 4.4 | 5.1 | 15.35 | 8.1 | 9 | 8.2 | 204 | 9.77 | 36.19 |
| 53b | SAK0244 | U | SF | 7 | 121 | 38 | 4.3 | 4 | 4.7 | 14.95 | 7.1 | 7.9 | 7 | 189 | 8.99 | 29.38 |
| 53b | SAK0245 | U | SF | 7 | 121 | 38 | 4.1 | 4.4 | 4.8 | 14.9 | 7 | 8.3 | NA | 279 | 9.50 | 29.63 |
| 09b | SAK0199 | NU | F | 8 | 75 | 0 | NA | NA | NA | NA | NA | 9.75 | NA | NA | NA | NA |
| 11 | SAK0192 | NU | SF | 8 | 72 | 53 | 4.9 | 4.8 | 5.4 | 18.05 | 10.5 | 8.6 | NA | NA | 6.98 | 39.25 |
| 11 | SAK0191 | NU | SF | 8 | 72 | 53 | 5 | 4.4 | 5.45 | 18.7 | 11.25 | 8.3 | 4.9 | NA | 6.32 | 38.68 |
| 31 | SAK0306 | NU | F | 8 | 78 | 0 | 5.45 | 5.1 | 5.5 | 17.6 | 11.7 | 9.25 | NA | 286 | 7.85 | 45.37 |
| 31 | SAK0307 | NU | F | 8 | 78 | 0 | 4.7 | 4.8 | 5.5 | 17 | 9 | 8.5 | 9.9 | NA | 7.68 | 38.02 |
| 47 | SAK0312 | NU | SF | 8 | 85 | 0 | 5.35 | 5.2 | 6.15 | 16.8 | 13.25 | 7.7 | 9.2 | 199 | 7.11 | 47.69 |
| 73 | SAK0363 | NU | F | 8 | 121 | 0 | 5.2 | 4.3 | 4.85 | 16.45 | 11.3 | 7.8 | NA | 220 | 7.48 | 37.37 |
| 1 | SAK0163 | U | SF | 8 | 51 | 13 | 5 | 4.6 | 6.3 | 17.9 | 14.1 | 11.1 | 8.9 | 316 | 9.14 | 42.80 |
| 5 | SAK0157 | U | F | 8 | 47 | 0 | 5.4 | 4.3 | 5.95 | 17 | 16.4 | 10.6 | NA | 367 | 9.58 | 43.47 |
| 05b | SAK0311 | U | F | 8 | 84 | 0 | 4.4 | 4.2 | 5.2 | 17.5 | 13.85 | 9.3 | 9 | 237 | 7.98 | 32.48 |
| 6 | SAK0168 | U | SF | 8 | 60 | 20 | 5.85 | 4.65 | 5.9 | 18.3 | 12.75 | 8.7 | NA | 269 | 6.89 | 48.47 |
| 6 | SAK0169 | U | SF | 8 | 60 | 20 | 5.35 | 3.55 | 5.8 | 17.6 | 13.75 | 9.4 | NA | 323 | 7.98 | 39.29 |
| 8 | SAK0153 | U | SF | 8 | 44 | 22 | 5.9 | 5.05 | 6 | 18.3 | 17.75 | 12.3 | 5.5 | NA | 9.74 | 51.20 |
| 8 | SAK0154 | U | SF | 8 | 44 | 22 | 5.7 | 5.45 | 6 | 17.9 | 15.6 | 13 | NA | NA | 10.71 | 51.26 |
| 9 | SAK0189 | U | F | 8 | 70 | 0 | 4.75 | 4.6 | 5.9 | 17.9 | 12.85 | 8.9 | 9.9 | NA | 7.33 | 39.17 |
| 9 | SAK0188 | U | F | 8 | 70 | 0 | 4.95 | 4.6 | 5.4 | 19.4 | 15 | 9.3 | NA | NA | 6.63 | 38.88 |
| 9 | SAK0187 | U | F | 8 | 70 | 0 | 5 | 5.1 | 5.4 | 19.55 | 16.4 | 10.7 | NA | NA | 7.52 | 41.23 |
| 30b | SAK0329 | U | F | 8 | 106 | 8 | 5.3 | 5.05 | 5.5 | 18.5 | 14.9 | 9.7 | 9.3 | 255 | 7.53 | 43.92 |
| 30b | SAK0330 | U | F | 8 | 106 | 8 | 5.05 | 4.9 | 5.7 | 17.5 | 12.3 | 9.3 | 7 | 282 | 7.98 | 42.04 |
| 35 | SAK0060 | U | SF | 8 | 65 | 26 | 5.3 | 5.1 | 5.3 | 18.3 | 15.6 | 10.9 | 11.4 | 250 | 8.63 | 43.29 |
| 35 | SAK0178 | U | SF | 8 | 65 | 26 | 5.3 | 4.4 | 5.3 | 18.2 | 14.5 | 10.4 | NA | NA | 8.32 | 40.38 |
| 38a | SAK0214 | U | F | 8 | 88 | 0 | 6.5 | 5.6 | 6.15 | 19.1 | 16.5 | 11 | 12.5 | 219 | 8.07 | 59.98 |
| 38a | SAK0217 | U | F | 8 | 88 | 0 | 6 | 5.25 | 6.4 | 19.45 | 14.5 | 10.4 | NA | NA | 7.38 | 54.90 |
| 42 | SAK0208 | U | F | 8 | 86 | 0 | 3.9 | 3.7 | 4.75 | 14.8 | 7.1 | 6.7 | NA | NA | 7.77 | 25.88 |
| 42 | SAK0207 | U | F | 8 | 86 | 0 | 4.3 | 3.9 | 4.7 | 16.2 | 11.95 | 7.4 | NA | NA | 7.29 | 29.04 |
| 42 | SAK0206 | U | F | 8 | 86 | 0 | 5 | 4.3 | 4.8 | 16.6 | 13.3 | 9.1 | 10 | NA | 8.58 | 35.74 |
| 48 | SAK0226 | U | F | 8 | 95 | 0 | 4.3 | 4.5 | 4.7 | 17.7 | 14.8 | 10.3 | 8.1 | 261 | 8.66 | 31.07 |
| 51 | SAK0211 | U | SF | 8 | 87 | 0 | 4.2 | 4.3 | 5 | 18.15 | 15.8 | 10.5 | NA | NA | 8.44 | 30.68 |
| 09b | SAK0197 | NU | F | 9 | 75 | 0 | 5.4 | 4.75 | 5.5 | 18.7 | 17.7 | 12.3 | 10 | 283 | 9.33 | 43.47 |
| 09b | SAK0198 | NU | F | 9 | 75 | 0 | 4.6 | 4.15 | 5.4 | 18.95 | 13 | 11.5 | NA | NA | 8.55 | 34.50 |
| 6 | SAK0167 | U | SF | 9 | 60 | 20 | 5.4 | 4.7 | 5.75 | 18 | 13.55 | 10.3 | NA | NA | 8.40 | 44.32 |
| 48 | SAK0225 | U | F | 9 | 95 | 0 | 4.1 | 4.5 | 4.6 | 18.4 | 15.3 | 11.4 | NA | NA | 8.94 | 29.30 |
| 51 | SAK0210 | U | SF | 9 | 87 | 0 | 5.3 | 4.8 | 5.2 | 17.5 | 16.3 | 11.1 | 7.9 | 289 | 9.52 | 41.63 |
| 51 | SAK0209 | U | SF | 9 | 87 | 0 | 5.5 | 4.5 | 5.5 | 17.4 | 16 | 11.3 | 9.6 | 315 | 9.79 | 43.20 |
| 13a | SAK0308 | NU | F | 10 | 78 | 0 | 5.7 | 5.3 | 6.25 | 19.75 | 17.15 | 13 | 12 | 322 | 8.98 | 51.71 |

Table S2. Total RNA reads, pre- and post-QC of nestling finch samples, and HISAT2 alignment rates to *Geospiza fortis* reference.

| Sample | Total Reads Pre-QC | Total Reads Post-QC | Alignment Rate |
| --- | --- | --- | --- |
| SAK0159_S75 | 26848858 | 25942013 | 91.8% |
| SAK0060_S23 | 18863623 | 18026570 | 86.0% |
| SAK0152_S25 | 22278342 | 21455609 | 87.8% |
| SAK0157_S39 | 22145819 | 21186414 | 86.0% |
| SAK0167_S48 | 23499497 | 22670471 | 86.9% |
| SAK0171_S26 | 19022442 | 18397526 | 86.5% |
| SAK0176_S54 | 19662231 | 18899534 | 84.1% |
| SAK0181_S22 | 20187913 | 19354348 | 85.8% |
| SAK0189_S19 | 19131752 | 18209251 | 85.7% |
| SAK0191_S18 | 21491783 | 20673627 | 85.3% |
| SAK0194_S33 | 21301972 | 20394515 | 85.0% |
| SAK0198_S17 | 26222486 | 25017039 | 86.3% |
| SAK0200_S16 | 28389046 | 26676144 | 85.6% |
| SAK0207_S8 | 21819109 | 20836487 | 84.7% |
| SAK0210_S42 | 19505710 | 18631390 | 85.5% |
| SAK0218_S56 | 26024025 | 25059514 | 87.4% |
| SAK0222_S3 | 22187688 | 21247944 | 85.4% |
| SAK0223_S2 | 24165607 | 22980306 | 84.2% |
| SAK0225_S36 | 21836556 | 20931210 | 87.7% |
| SAK0245_S29 | 19343382 | 18606756 | 87.3% |
| SAK0247_S53 | 21698902 | 20841716 | 87.4% |
| SAK0250_S51 | 20416550 | 19622930 | 87.1% |
| SAK0304_S58 | 21614650 | 20791300 | 86.0% |
| SAK0307_S59 | 21184654 | 20383881 | 87.2% |
| SAK0308_S57 | 17607026 | 16772670 | 85.1% |
| SAK0311_S11 | 26553445 | 25405743 | 84.0% |
| SAK0313_S14 | 25277694 | 24240886 | 86.6% |
| SAK0317_S6 | 26843299 | 25689072 | 86.0% |
| SAK0326_S47 | 21971053 | 19799958 | 86.6% |
| SAK0328_S41 | 26248067 | 24532889 | 86.7% |
| SAK0330_S65 | 17186825 | 16409058 | 85.4% |
| SAK0331_S52 | 19901554 | 19132563 | 86.8% |
| SAK0333_S55 | 20280118 | 19506047 | 85.9% |
| SAK0341_S50 | 18030947 | 17293197 | 87.7% |
| SAK0347_S70 | 18440662 | 17537461 | 80.7% |
| SAK0352_S64 | 21398182 | 20582596 | 87.1% |
| SAK0354_S38 | 16705859 | 15863427 | 87.0% |
| SAK0358_S60 | 21481516 | 20690347 | 86.0% |
| SAK0362_S34 | 20234296 | 19374380 | 84.4% |
| SAK0364_S20 | 43727416 | 42156991 | 86.3% |
| SAK0367_S9 | 23041035 | 22026003 | 86.0% |
| SAK0612_S46 | 19764787 | 18900722 | 87.0% |
